# Biased signaling at NTSR1 differentially regulates inhibitory synaptic transmission in the extended amygdala and suppresses motivated feeding in mice

**DOI:** 10.64898/2026.04.30.722083

**Authors:** Sarah E. Sizer, Alex R. Brown, Josephine K. Anderson, Abigail E. Summerlin, Isabella Girgis, Stephen Y. Yang, Madigan L. Bedard, Steven H. Olson, Lauren M. Slosky, Gina M. Leinninger, Zoé A. McElligott

**Affiliations:** Bowles Center for Alcohol Studies, UNC Chapel Hill, Chapel Hill, NC, USA; Department of Pharmacology, UNC Chapel Hill, Chapel Hill, NC, USA; Sanford Burnham Prebys, La Jolla, USA; Department of Pharmacology, University of Minnesota Medical School, Minneapolis, USA; Department of Physiology, Michigan State University, East Lansing, MI, USA; Department of Psychiatry, UNC Chapel Hill, Chapel Hill, NC, USA

**Author notes:** These authors contributed equally to the project (Sarah E. Sizer, Alex R. Brown).

**Keywords:** Neurotensin, biased modulation, GABA, central nucleus of the amygdala, bed nucleus of the stria terminalis

## Abstract

Maladaptive consummatory behaviors can arise from dysregulated circuits, like those in the extended amygdala, governing motivation and feeding behaviors. Neurotensin (NTS), expressed throughout the central, peripheral, and enteric nervous systems, has well-established roles in energy balance and feeding. SBI-553, a biased allosteric modulator of NTSR1, recruits β-arrestin while attenuating Gq-mediated signaling. We used SBI-553 to examine NTS modulation GABAergic signaling in the central amygdala (CeA) and bed nucleus of the stria terminalis (BNST) and probed its effects on food consumption in mice. We found that NTS and SBI-553 differentially modulate GABAergic neurotransmission across extended amygdala subregions. *In vivo*, SBI-553 reduces palatable food consumption in both fed and food-deprived mice, with greater reductions under fasted conditions, altering activation across CeA subregions in a sex- and feeding-state-dependent manner. In the BNST, differences in SBI-553 altered cFos activity were observed between feeding states.

**Graphical Abstract:** 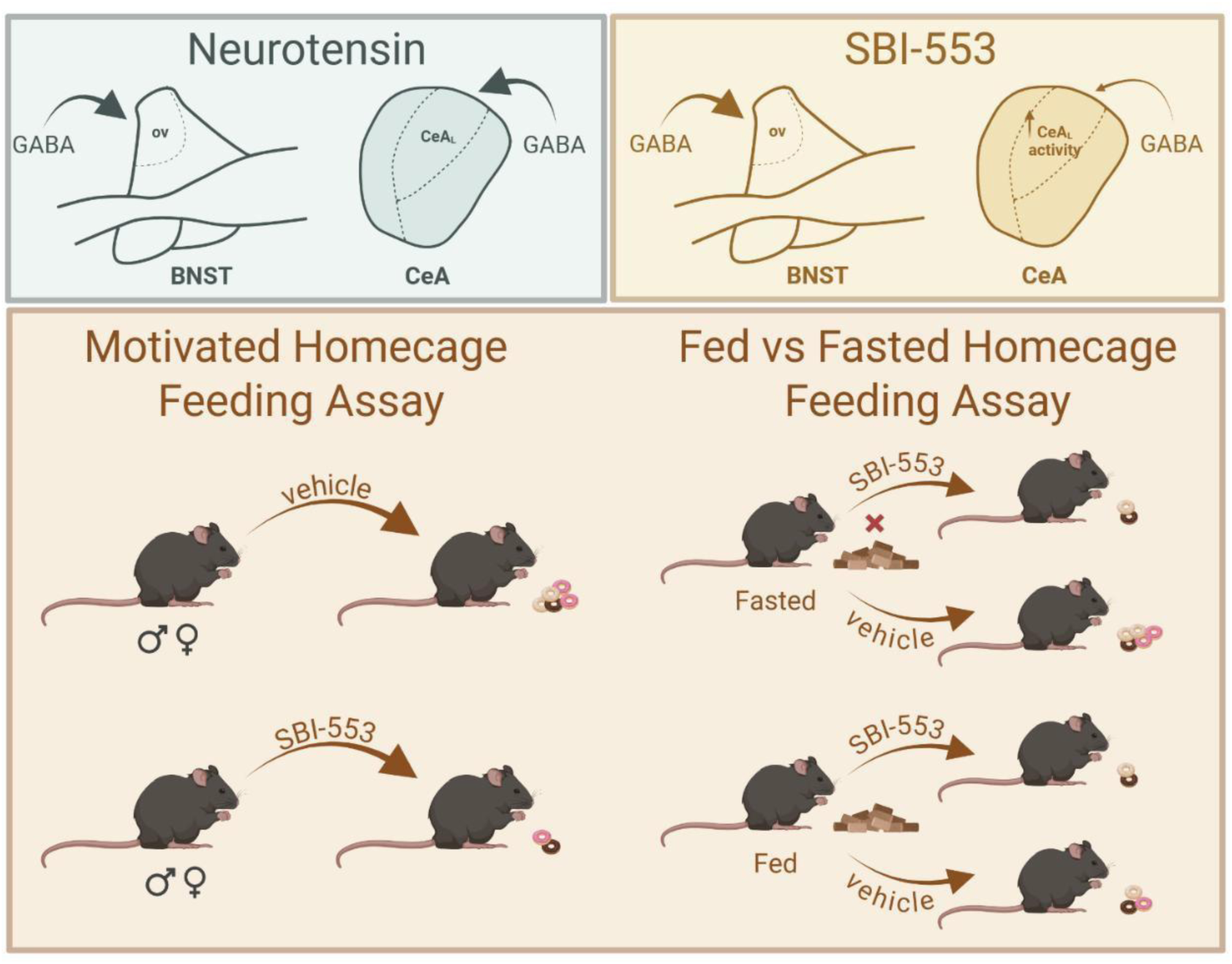

**Highlights:**

- NTS enhances GABAergic transmission in the CeAL and the ovBNST
- SBI-553 blocks NTS-induced increases in sIPSC frequency in the CeAL but not in the ovBNST
- SBI-553 attenuates the feeding of a palatable high-carbohydrate food
- The effect of SBI-553 on feeding is driven by energy deficit/motivation to feed

## Introduction

Neurotensin (NTS) is a 13-amino acid peptide expressed throughout the brain, enteric nervous system, reproductive system, and periphery that modulates feeding behavior and energy homeostasis. Aberrant NTS signaling is associated with the pathogenesis of body weight and feeding disorders [1,2], and clinically obese individuals, insulin-resistant individuals, and individuals with obesity-related genetic conditions have elevated plasma levels of NTS or its precursor, pro-NTS [3–5]. Elevations in plasma pro-NTS are correlated with a higher risk of developing obesity, type 2 diabetes, and cardiovascular disease [5]. Preclinical studies complement clinical research, suggesting that peripheral NTS promotes fat absorption and weight gain, yet the neurophysiological effects of NTS signaling on consummatory neurocircuits and its influence on feeding behavior remain less clear. While early research suggested that NTS had opposing effects in the brain versus the periphery and promoted weight loss, the advent of the Cre-lox system has enabled more precise isolation and manipulation of NTS circuits [6–11]. This recent body of work suggests that the effects of NTS on synaptic plasticity and behavior are not exclusively anorectic, but instead vary across brain regions [12,13]. The co-occurrence of neuropeptides, small neurotransmitters, and signaling molecules within these distinct circuits further complicates the effects of the NTS system on feeding behavior [13–15].

The central amygdala (CeA) is a sexually dimorphic brain region in the extended amygdala involved in the expression of fear, anxiety, pain, reward-seeking, and feeding behaviors [16,17]. Composed of GABAergic neurons, the lateral (CeAL), medial (CeAM), and capsular (CeAC) subregions are highly interconnected, yet serve distinct functions in orchestrating CeA output [18]. The distinct subregions also receive differential inputs affecting neuronal activity [19–23]. Although the CeAM is generally regarded as the primary output structure of the CeA [24–26], we have shown that the CeAL also sends dense efferent projections to regions like the PBN [13]. Therefore, the high degree of interconnectivity between inter- and extra-amygdalar circuits complicates our understanding of how information flows through the CeA to encode behavior [13,25,27,28]. While present in both the CeAL and CeAM, CeA^NTS^ are most abundant in the CeAL, and we and others have found that CeA^NTS^ neurons co-express several other markers, including somatostatin, corticotropin-releasing factor, and PKCδ [13,19,29,30]. CeA^NTS^ neurons regulate fear learning, spatial learning, reward, and sleep behaviors [31,32]. Furthermore, previous work from multiple laboratories including our own has shown that CeA^NTS^ projections to the PBN promote the consumption of alcohol, food, and sweet fluids [13,19,28,32]. Additionally, we demonstrated that blunting GABA release from CeA^NTS^ neurons reduces alcohol consumption in male mice and reduces thermal nociception in female mice [33,34]. The bed nucleus of the stria terminalis (BNST) is another component of the extended amygdala and is homologous to the CeA with similarities in GABAergic cellular composition, protein and neuropeptide expression, and afferent and efferent projections [35–37]. While the BNST is broadly divided by the anterior commissure into ventral (vBNST) and dorsal (dBNST) subregions, it is further divided into multiple subnuclei [38,39]. The oval nucleus (ovBNST) is the subnucleus most analogous to the CeAL, in part because it contains the highest expression of NTS (BNST^NTS^) neurons [40]. Although ovBNST neuropeptide co-expression patterns vary by species, BNST^NTS^ neurons overlap significantly with BNST^CRF^ and BNST^pdyn^ populations and project to the LH, PAG, PBN, and VTA to mediate stress- and feeding-related behaviors [40–49]. oVBNST neurons are sensitive to changes in metabolic state. For example, caloric state and binge-like sucrose intake alter the synaptic plasticity at ovBNST GABAergic synapses via endocannabinoid, dopaminergic, and putative neurotensinergic mechanisms [44,50]. Since release of NTS by BNST^NTS^ neurons also upregulates CeA GABAergic inputs [40], this circuit may be uniquely positioned to regulate feeding behavior, suggesting that targeting the NTS system could be beneficial for treating body weight and feeding disorders.

NTS binds to two G-protein-coupled receptors (GPCRs), the high-affinity neurotensin 1 receptor (NTSR1) and the low-affinity neurotensin 2 receptor (NTSR2) [51–53]. Global knockouts (KOs) of NTSR1 suggest that NTSR1 is the site of the anorectic action of NTS., however not all generated KOs demonstrate increased bodyweight suggesting that other genetic differences contribute [54]. NTSR1 canonically signals through a Gαq-dependent mechanism, but can also recruit β-arrestin, which may subsequently promote receptor internalization and mediate signaling [55–57]. However, NTSR1 is highly promiscuous and can also signal through Gαs, Gαi/o, Gα11, Gα12, Gα13, and Gα15 [58–60]. Although drugs targeting NTS receptors show therapeutic potential across several pathologies, traditional NTS receptor agonists cause undesirable side effects, such as sedation, hypotension, and hypothermia [55,61–63]. NTSR1-biased allosteric modulators (BAMs) could mitigate these side effects by shifting the relative signaling contributions of G-proteins and β-arrestin. SBI-553 is a NTSR1-BAM that antagonizes NTSR1 Gαq-mediated signaling in favor of the β-arrestin pathway, without impacting NTSR2-mediated signaling [64,65]. Previous findings from our laboratory and others show that SBI-553 reduces alcohol intake, alcohol-related behaviors and physiology, psychostimulant self-administration, and expression of reward learning in rodents [33,55] without altering locomotor, avoidance, or memory behavior, suggesting its utility as a pharmacotherapeutic for substance use disorders. Interestingly, our previous study also showed that SBI-553 reduces post-test refeeding following the novelty-suppressed feeding test (NSFT) [33], prompting additional questions about its effects on motivated feeding behavior. Based on this data and the well-established roles of the BNST and CeA in feeding behaviors, the present study used a combination of whole-cell patch clamp electrophysiology, cFos immunohistochemistry, and behavioral assays to understand the neurophysiological effects of SBI-553 on consummatory neural nodes and its influence on feeding behavior. Our *ex vivo* electrophysiology data suggest that SBI-553 differentially alters NTS modulation of GABAergic synaptic transmission within the CeA and BNST. Our *in vivo* findings suggest that systemic injection of SBI-553 exerts sex- and feeding state-dependent effects on palatable food intake. Together, these data suggest that SBI-553 may be a promising pharmacotherapeutic for hedonic feeding dysregulation.

## Methods

### Subjects, housing conditions, and feeding details

All procedures were conducted in accordance with the NIH *Guide to Care and Use of Laboratory Animals* with approval from the Institutional Animal Care and Use Committee at UNC-Chapel Hill. All animals were maintained under continuous veterinary care from the Division of Comparative Medicine. Male and female mice were used for all experiments and are represented in the results with distinct data symbols.

C57/BL6J mice (>8 weeks old; Jackson Laboratories) were used for *ex vivo* electrophysiology and *in vivo* naïve animal studies (Immunofluorescence, see below). Mice were housed in normal light conditions (7AM light ON-7PM light OFF) and allowed to acclimate to the vivarium for at least one week prior to any experiments. All C57 mice received *ad libitum* access to chow (5V5R PicoLab^®^ Select Rodent Diet 50 IF/6, LabDiet) and water.

Nts-Cre mice (8-35 weeks old, bred in house) were used in *in vivo* experiments investigating the role of SBI-553 on feeding behavior and cFos immunoreactivity. Our previous investigation of SBI-553 [33] used C57BL6/J mice; however, in this study, Nts-Cre animals were selected for two reasons: (1) Test if SBI-553 administration would similarly alter consummatory behavior across mouse strains (NTS-cre mice are on a mixed 129/C57 background); (2) Allow manipulation of NTS neurocircuitry using Cre-dependent viral approaches in future studies. Additionally, to match our previous study and to investigate feeding behavior during the active cycle, animals were housed in groups of up to 5 per cage and maintained on a reverse light cycle (7AM light OFF-7PM light ON) for at least one week prior to the start of the experiment, after which they were all single housed until completion. All animals had access to *ad libitum* water during all periods, and *ad libitum* chow (5V5R PicoLab^®^ Select Rodent Diet 50 IF/6, LabDiet) except during the food restriction period. Additionally, Froot Loops (FL; Kellogg’s) were used as a highly palatable food. See Supplemental Table 2 for a nutritional comparison of the foods.

#### Whole cell patch clamp electrophysiology

Male and female C57/BL6J mice (9-22 weeks old) were anesthetized with isoflurane, decapitated, and their brains were rapidly extracted. Brains were submersed in oxygenated, ice-cold high-sucrose low Na^+^ artificial cerebral spinal fluid (aCSF; in mM: 194 sucrose, 20 NaCl, 4.4 KCl, 2 CaCl2, 1 MgCl2, 1.2 NaH2PO4, 10 glucose, 26 NaHCO3) and sliced at 250 μM thickness with a Leica VT1000S vibratome. Coronal slices containing the CeA and BNST were placed in a holding chamber containing standard aCSF (in mM: 124 NaCl, 4.4 KCl, 2 CaCl2, 1.2 MgSO4, 1 NaH2PO4, 10 glucose, 26 NaHCO3; osmolarity ∼300 mOsm), heated to 32°C for at least 1 hour prior to recording. Slices were transferred to the recording chamber, where aCSF containing 3 mM kynurenic acid (glutamate receptor antagonist) was heated (30°C) and continuously perfused at a rate of 2 mL/minute. Patch electrodes (3-6 MΩ) were pulled using a micropipette puller (P-97 Sutter Instruments) and filled with cesium-chloride internal solution (in mM: 130 CsCl2, 38 EGTA, 10 HEPES, 2 ATP, 2 GTP, 1 QX-314; pH ∼7.35, osmolarity ∼285 mOsm). Neurons were held at −80 mV to record the frequency, amplitude, area, and kinetics of spontaneous inhibitory postsynaptic currents (sIPSCs) during a 5-minute baseline and during a 10-minute application of neurotensin (1 µM) and/or SBI-553 (10 µM). A subset of experiments concluded with a 5-minute drug washout. 400 µM neurotensin (diluted in double-distilled H2O) and 2 mM SBI-553 (diluted in 5% hydroxypropyl-β-Cyclodextrin or 1.67% DMSO) stock solutions were stored at −20°C and diluted in aCSF containing kynurenic acid. Electrophysiology recordings with a change in access resistance >20% were excluded from analysis. All electrophysiology data were analyzed using Clampfit 11 (Molecular Devices) or Easy Electrophysiology (Easy Electrophysiology, Ltd.). Cellular properties are reported in Supplemental Table 1.

#### Naive Animal SBI-553 Injections

16 male and 16 female C57/BL6J mice (food/water *ad. libitum*) were injected with either an SBI-553 (12 mg/kg I.P.) or equivolume vehicle (10 mL/kg, I.P.). For each sex, 8 animals were injected with SBI-553 (12 mg/kg), and 8 were injected with 5% hydroxypropyl-β-Cyclodextrin (vehicle). After 90 minutes, mice were transcardially perfused with 4% paraformaldeyde, and brains were stored in 30% sucrose at 4°C for cFos immunohistochemistry.

#### Immunofluorescence cFos staining

Perfused brains were sliced using a Leica cryostat (40 μm thick) and stored in 50/50 PBS/Glycerol at 4°C. Slices containing the BNST and the CeA were stained for the immediate early gene cFos. Fixed coronal slices were rinsed twice in 0.01 M PBS (Fisher Bioreagents) for 10 minutes. Slices were incubated for 20 minutes in a 3% hydrogen peroxide (30% H2O2)-PBS solution. Following two PBS washes, slices were incubated in a blocking buffer (0.3% Triton X-100 (PBS-T solution) and 1% Bovine Serum Albumin (BSA) for one hour. Slices were incubated for 48 hours at 4°C in rabbit anti-C-FOS primary antibody (1:3,000; Synaptic Systems, Germany) diluted in PBS-T and 0.5% BSA, except for a subset of no-primary controls. Slices were washed three times with a TNT wash for 10 minutes (stored at 4°C) (TNT wash: 0.1 M TRIS HCl, 0.15 M NaCl, 0.5% Triton X-100) and then washed in a TNB buffer for 30 minutes. Slices were incubated with HRP-conjugated goat anti-rabbit (1:200) secondary antibody diluted in TNB buffer for one hour. Slices were washed with TNT buffer and then incubated in Cy3 (1:50) diluted in TSA amplification diluents. Slices were washed twice with TNT buffer and incubated with DAPI diluted in PBS (1:1000) for 5 minutes. Slices were washed with TNT buffer and rinsed with PBS before mounting with Vector Vibrance mounting medium without DAPI (ThermoFisher). Images were taken with an ECHO confocal microscope or the VS200 Slide Scanner provided by the UNC Hooker Imaging Core. The number of cFos-positive cells in the lateral, medial, and capsular subregions of the CeA and the oval, dorsal and ventral subregions of the BNST were quantified using QuPath Software [66].

#### Feeding Assays

Motivated homecage feeding assay (MHFA): 24 male and 18 female Nts-Cre mice [15] (no surgical/viral manipulation) underwent the Motivated Homecage Feeding Assay (MHFA). On day 1, animals were singly housed. On day 2, mice were given access to 2 Froot Loops (FL) in a plastic weigh boat to acclimate them to the novel food. On day 3, cages were thoroughly checked for FL consumption, and animals were given another 2 FL in the morning (∼3 hours into dark cycle). At least 4 hours after providing the FL, cages were checked for FL consumption, and all chow was removed to start food deprivation. On day 4, Test Day, ∼20 hours after chow removal, animals received an injection of SBI-553 (12 mg/kg, I.P.) or vehicle (5% Hydroxypropyl β-cyclodextrin, 10 mL/kg, I.P.) 30 minutes prior to receiving a weigh boat containing 10 FL. After 10 minutes of FL access, bedding was checked for crumbs/partially eaten FL, and any discovered FL was added back to the weigh boat. The difference in the weigh boat mass was quantified as consumption and reported as kcal consumed per kg of animal body weight (kcal/kg). 1 hour following the removal of the weigh boat, we trascardially perfused a subset of these animals for immunofluorescence cFos staining (see methods above). Mice that failed to consume any amount of FL on either the acclimation days or the test day were excluded from analysis (n=3 male mice).

Fasted vs fed homecage feeding assay (FvFHFA): A separate cohort of 15 male and 11 female Nts-Cre mice (no surgical/viral manipulations) underwent a similar experiment to assess the impact of hunger state on SBI-553-induced changes in feeding. FvFHFA was identical to MHFA with two notable exceptions: (1) half of the animals (8 males, 5 females) were assigned to the fasted group and were food-deprived on day 3, allowing for the assessment of fasting on consummatory behaviors. The other half (7 males, 6 females) were assigned to the fed group, which had food *ad libitum*. (2) After the completion of the Test Day (Day 4), animals had chow replaced and were allowed to re-establish baseline consummatory behaviors for the following 3 days. Then, the 4-day test was repeated with the second treatment (SBI-553 or Vehicle) to measure within-animal treatment efficacy. The order of vehicle and SBI-553 treatment was counterbalanced across weeks, and we did not observe an effect of injection order (Fig 7D). After the second treatment session, we transcardially perfused a subset of these animals 1 hour after food removal for immunofluorescence cFos staining. Mice that failed to consume any amount of FL during the acclimation period were excluded from analysis (n=1 male mouse).

#### Data Analysis

All statistical analyses were performed in GraphPad Prism (version 10.6.1 and 11.0.2). Significance was set at *p*≤0.05, and effects at *p*<0.1 were considered trends toward significance. Franklin and Paxinos, *The Mouse Brain in Stereotaxic* Coordinates, 3^rd^ edition was used to reference brain regions. Due to well-established sex differences in feeding behavior, within the NTS system [67,68], and in the physiology and morphology of the BNST and CeA, data were stratified by sex for all initial analyses. In some datasets where no sex differences were observed, data were collapsed across sex to increase statistical power; these instances are noted in the results. Data were analyzed using unpaired Student’s T-test two-tailed), Friedman test (cumulative distributions), regular two-way ANOVAs, repeated measures 2-way ANOVAs, or repeated measures 3-way ANOVAs designs, where appropriate. Following a significant main effect or interaction in the ANOVAs, post-hoc comparisons were conducted using Bonferroni correction for designs with two levels per factor or Tukey’s test for designs with more than two levels per factor. This allowed us to determine whether one group drives a main effect or interaction.

Specific statistical tests and sample sizes are reported in the results. For effect size calculation *η^2^P* = SSeffect/(SSeffect +SSerror) and *ω^2^P* = [*df*effect * (MSeffect – MSerror)]/ [*df*effect * MSeffect + (N – *df*effect) x MSerror] were calculated from the ANOVA, where generally cut off values for small effect sizes are 0.01, medium are 0.06, and large are 0.14 [69].

## Results

Prior research in the extended amygdala has focused on NTS modulation of GABAergic signaling in the oval BNST (ovBNST), where NTS mediates a presynaptically expressed inhibitory long-term potentiation (iLTP) [70] and enhances intra-CeA GABAergic signaling [71]. We and others have previously reported that bath application of, or depolarization induced endogenous NTS release results in an iLTP of electrically evoked GABAergic release in the BNST of rats and mice, and that endogenous release produces a similar adaptation in the CeA→BNST pathway in mice mediated by NTSR1 [42,68]. The Dumont lab further explored the effects of this depolarization on excitatory transmission in the ovBNST and found that the depolarization inhibited excitatory transmission. In ∼90% of the cells recorded this effect was blocked by the opioid antagonist naloxone [73], suggesting that NTS selectively modulates inhibitory transmission. Furthermore, selective NTSR2 agonists inhibited evoked IPSCs [74]. Ergo, we sought to understand whether NTS modulates spontaneous GABAergic signaling across CeA and BNST subregions without releasing other factors via electrical stimulation or neuronal depolarization. We used whole-cell patch clamp electrophysiology to record spontaneous inhibitory postsynaptic currents (sIPSCs) in naïve male and female mice at baseline, during NTS (1 µM) perfusion, and after a drug washout.

NTS significantly increases the average sIPSC frequency onto neurons within the lateral CeA (CeAL), but surprisingly and unlike our previous observations with evoked release, the elevation in sIPSC frequency returns to baseline during the washout (Fig. 1B; two-way repeated measures ANOVA; males: n= 8 cells, 5 mice; females: n= 8 cells, 6 mice; significant main effect of NTS *F*(1.26,17.56)=12.94, η^2^p= 0.480, ω^2^p=0.599, *p*=0.0012; no main effect of sex *F*(1,14)=1.327, *p*=0.269, and no NTS x sex interaction *F*(2,28)=0.568, *p*=0.573; Tukey’s multiple comparisons test; baseline vs. NTS, *p*=0.0033, NTS vs. washout, *p*=0.0094). The cumulative distributions show similar leftward shifts in the interevent interval following NTS perfusion (Sup Fig. 1G). In the CeAL, NTS does not alter the sIPSC amplitude (Figure 1C; two-way repeated measures ANOVA; no main effect of NTS *F*(1.38,19.33)=0.889, *p*=0.391; no main effect of sex *F*(1,14)=0.214, *p*=0.651, and no NTS x sex interaction *F*(2,28)=0.596, *p*=0.502) or the sIPSC area (Fig. 1D; two-way repeated measures ANOVA; no main effect of NTS *F*(1.288,18.04)=2.349, *p*=0.138; no main effect of sex *F*(1,14)=1.406, *p*=0.255, and no NTS x sex interaction *F*(2,28)=0.495, *p*=0.615). We did not observe changes in the sIPSC kinetics (Sup Fig. 1A,B). These findings, along with prior reports [40,71], suggest that NTS upregulates GABAergic input onto CeAL neurons in males and females, likely via a presynaptic mechanism.

**Figure 1.**
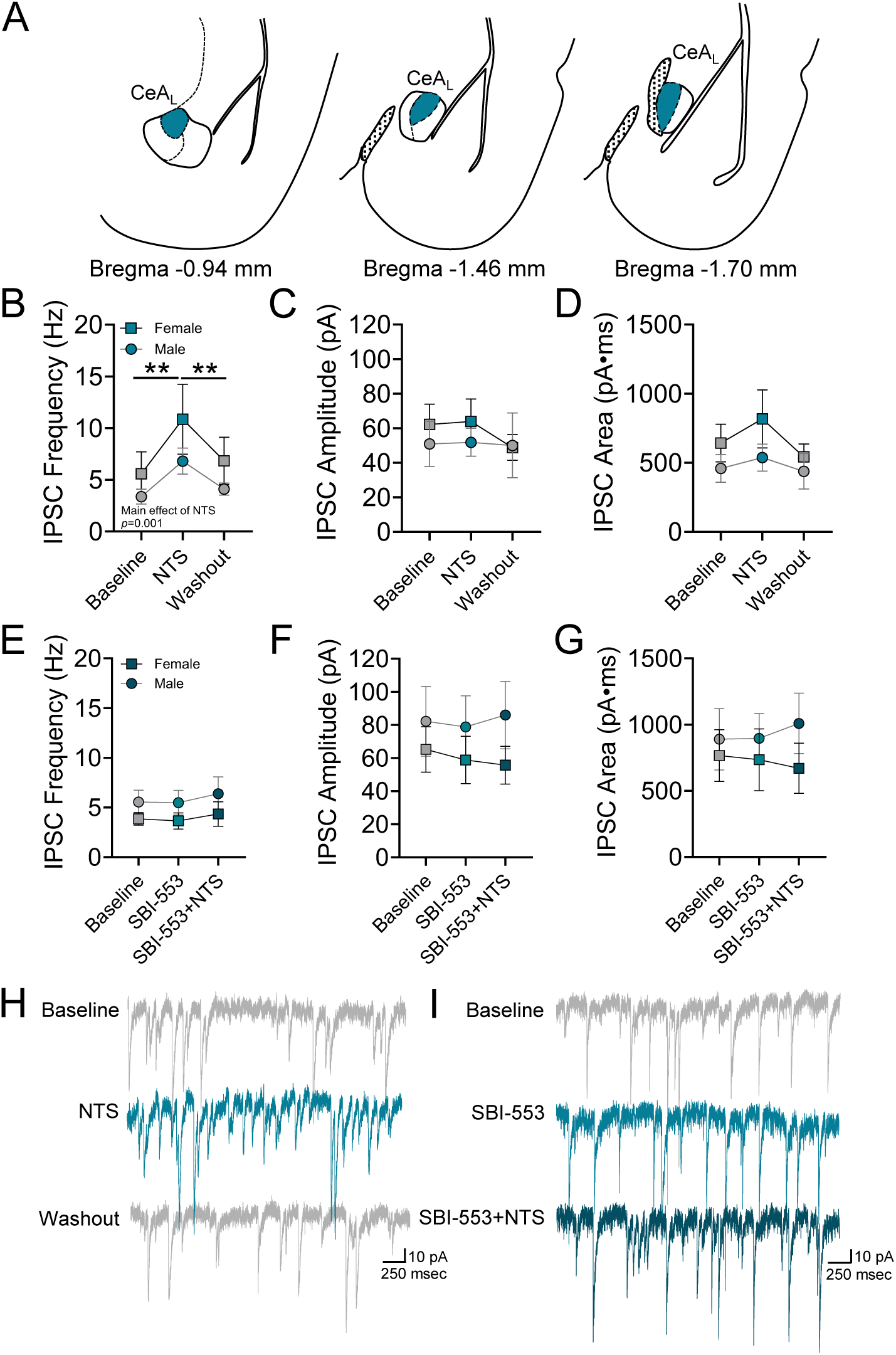
SBI-553 blocks NTS-induced increases in GABAergic neurotransmission onto CeA_L_ neurons independently of sex **(A)** Schematic of CeA subregions across the AP axis and recording location of electrophysiology experiments denoting all recordings were in the CeA_L_ in this figure (teal) **(B)** Neurotensin increases the sIPSC frequency in both males and females. This effect returns to baseline levels during the washout. Neurotensin did not change the **(C)** IPSC amplitude or **(D)** IPSC area in either sex **(E)** SBI-553 alone does not alter the IPSC frequency, but blocks NTS from increasing IPSC frequency. SBI-553 alone of in combination with NTS does not alter **(F)** IPSC amplitude or **(G)** IPSC area. Representative traces of **(H)** NTS perfusion and **(I)** SBI-553 and NTS perfusion. ** p<0.01

SBI-553 is a BAM that antagonizes NTSR1 Gq-mediated signaling in favor of the β-arrestin pathway [55,60]. We used whole-cell patch clamp electrophysiology to determine whether SBI-553, alone or in combination with NTS, modulates GABAergic neurotransmission in the CeAL. While SBI-553 (10 µM) alone does not alter the sIPSC frequency, pre-treatment with SBI-553 occludes NTS-induced increases in sIPSC frequency in both sexes (Fig. 1E; two-way repeated measures ANOVA; males: n= 7 cells, 5 mice; females: n= 7 cells, 6 mice; no main effect of drug wash on *F*(1.232,14.78)=0.979, *p*=0.358; no main effect of sex *F*(1,12)=1.511, *p*=0.243, and no drug x sex interaction *F*(2,24)=0.041, *p*=0.887). The cumulative distributions show that SBI-553 attenuates the leftward shifts in the interevent interval with NTS alone (Sup Fig. 1H). SBI-553 alone and in combination with NTS does not alter the sIPSC amplitude (Fig. 1F; two-way repeated measures ANOVA; no main effect of drug wash on *F*(1.729,20.75)=0.440, *p*=0.621; no main effect of sex *F*(1,12)=0.930, *p*=0.354, and no drug x sex interaction *F*(2,24)=0.883, *p*=0.426) or the sIPSC area (Fig. 1G; two-way repeated measures ANOVA; no main effect of drug *F*(1.798,21.57)=0.049, *p*=0.940; no main effect of sex *F*(1,12)=0.529, *p*=0.481, and no drug x sex interaction *F*(2,24)=1.080, *p*=0.356). These findings suggest that SBI-553 prevents NTS-induced increases in GABAergic input by attenuating NTSR1 Gq-mediated signaling.

Since NTS^+^ neurons are also present, although less abundant, in the medial CeA (CeA_M_) [13], we similarly recorded CeA_M_ neurons after bath application of NTS. Akin to previous reports in rats, females exhibit a higher sIPSC frequency than males in the CeA_M_ [75,76]. NTS application, however, did not significantly change the sIPSC frequency in either sex (Fig. 2B; two-way repeated measures ANOVA; males: n= 7 cells, 7 mice; females: n= 8 cells, 6 mice; no main effect of NTS *F*_(1.575,20.47)_=1.868, *p*=0.185; a significant main effect of sex *F*_(1,13)_=6.253, *p*=0.027, η^2^ = 0.598, ω^2^ =0.715, and no NTS x sex interaction *F* =1.167, *p*=0.327), or shift the sIPSC frequency cumulative distributions (Sup Fig. 1I). NTS also did not alter the sIPSC amplitude (Fig. 2C; two-way repeated measures ANOVA; no main effect of NTS *F*_(1.547,20.11)_=0.251, *p*=0.724; main effect of sex *F*_(1,13)_=2.181, *p*=0.164, and no NTS x sex interaction *F*_(2,26)_=1.760, *p*=0.192) or the sIPSC area (Fig. 2D; two-way repeated measures ANOVA; no main effect of NTS *F*_(1.935,25.15)_=0.969, *p*=0.391; main effect of sex *F*_(1,13)_=2.621, *p*=0.129, and no NTS x sex interaction *F*_(2,26)_=0.462, *p*=0.635). NTS does not significantly alter the sIPSC kinetics, although there was a trend for a main effect of NTS on IPSC decay (*p*=0.053, Sup Fig. 1C-D). These findings suggest that NTS exerts region-specific effects within the CeA, with the most robust effects occurring in the CeA_L_, likely due to the anatomical distribution of NTS-expressing cells. More broadly, these findings add to a growing body of work describing region- and sex-dependent differences in synaptic plasticity within the CeA [23,34,75–78].

**Figure 2.**
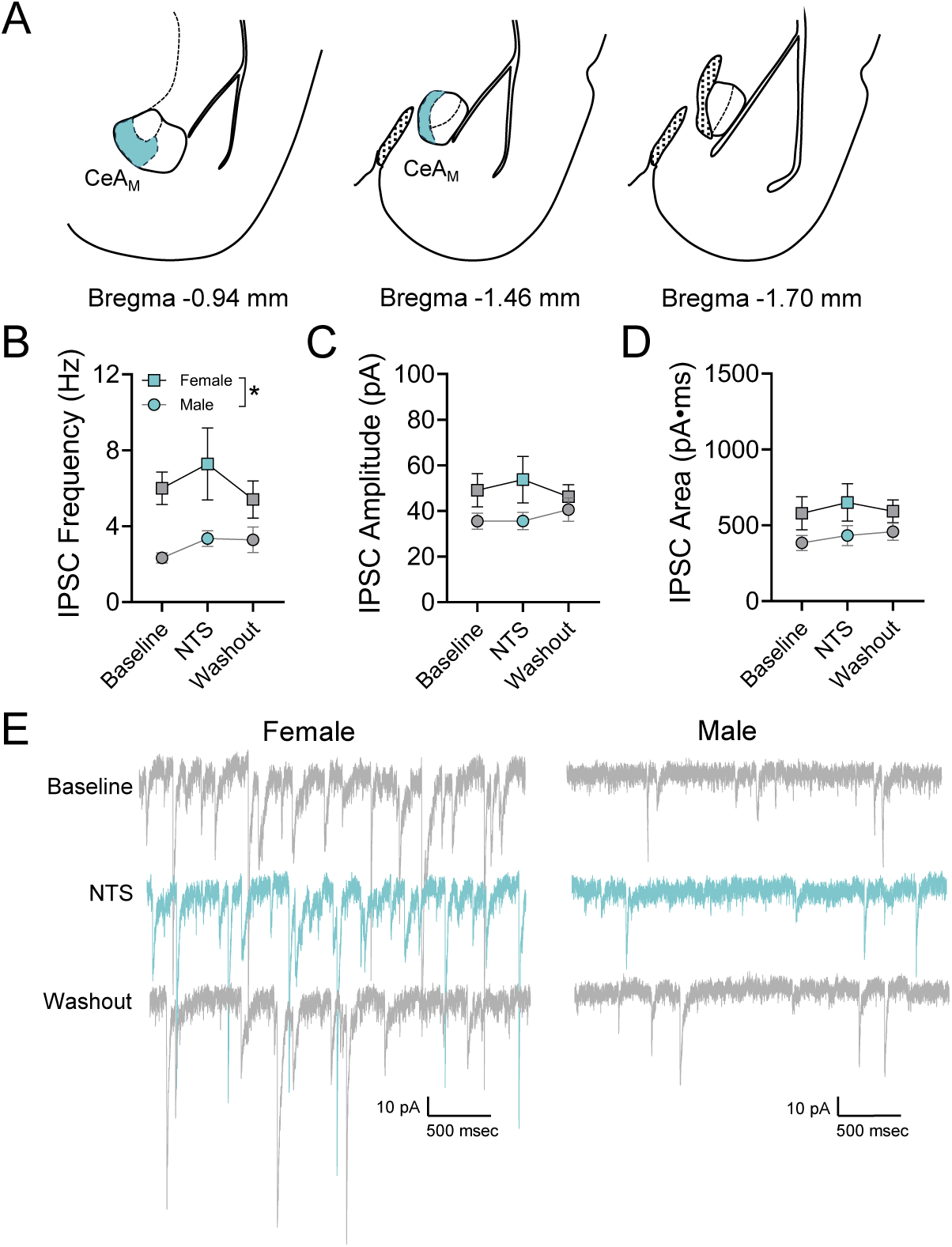
Neurotensin does not alter GABAergic neurotransmission onto CeA_M_ neurons **(A)** Schematic of CeA subregions across the AP axis and recording location of electrophysiology experiments denoting all recordings were in the CeA_M_ (light blue) in this figure **(B)** Females have a higher IPSC frequency at baseline than males, but neurotensin does not alter the sIPSC frequency in either sex. Neurotensin also does not alter the **(C)** IPSC amplitude or **(D)** IPSC area. Representative traces of NTS perfusion in **(E)** females and males. * p<0.05

In the ovBNST, endogenous NTS upregulates GABAergic input, including that from GABA inputs from CeA projections in male mice and rats [40,72], yet it remains unclear whether NTS promotes sex-specific effects on ovBNST synaptic transmission. We found NTS increases the sIPSC frequency and shifts the cumulative distributions leftwards in both sexes (Fig. 3C, Sup Fig. 1J; two-way repeated measures ANOVA; males: n= 8 cells, 6 mice; females: n= 10 cells, 6 mice; significant main effect of NTS *F*_(1.416,22.65)_=8.823, *p*=0.003, η^2^_p_= 0.335, ω^2^_p_= 0.465; no main effect of sex *F*_(1,16)_=0.007, *p*=0.936, and no NTS x sex interaction *F*_(2,32)_=0.804, *p*=0.457; Tukey’s multiple comparisons test; baseline vs. NTS, *p*=0.027, NTS vs. washout, *p*=0.0084). In contrast to our findings in the CeA_L_, however, NTS promoted a significant sex x amplitude interaction in sIPSC amplitude (Fig. 3D; two-way repeated measures ANOVA; no main effect of NTS *F*_(1.384,22.14)_=0.583, *p*=0.506; significant main effect of sex *F*_(1,16)_=5.419, *p*=0.033, η^2^_p_= 0.584, ω^2^_p_= 0.709; and, NTS x sex interaction *F*_(2,32)_=3.904, *p*=0.049, η^2^_p_= 0.196, ω^2^_p_= 0.244, Bonferroni’s multiple comparisons test shows a significant difference in response to NTS, p=0.030, between male and female mice), demonstrating that NTS promotes sex-specific effects on GABAergic signaling in the ovBNST. While NTS similarly increase sIPSC area, the effect was not statistically significant (Fig 3E, no main effect of NTS F_(1.405, 22.47)_=0.498, p=0.5496, no main effect of sex F_(1,16)_=2.970, p=0.104; no NTS x sex interaction F_(2,32)_=0.1836). These findings may suggest that the changes in frequency and amplitude are occurring on different synapses in the male mice. In contrast to the CeA, we found NTS changes the sIPSC rise time kinetics, but not the decay (Sup Fig. 1E, significant main effect of NTS F_(1.919, 30.70)_=4.760, p=0.017; no effect of sex F_(1,16)_=0.502, p=0.489; no NTS x sex interaction F_(2,32)_=0.180; p=0.828).

**Figure 3.**
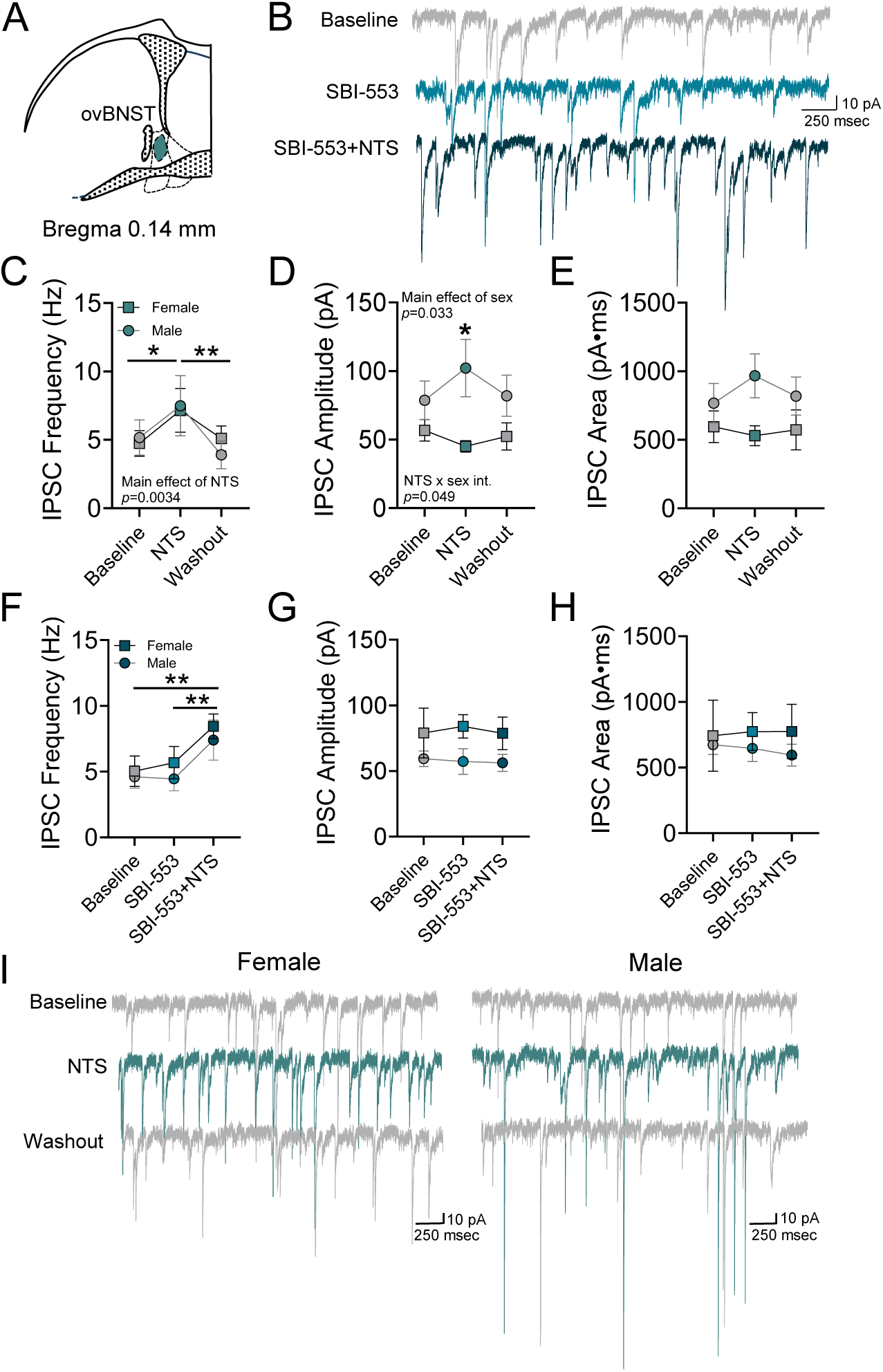
SBI-553 has complex actions on inhibitory transmission onto ovBNST neurons **(A)** Schematic of the ovBNST subnuclei and recording location of electrophysiology experiments (teal) **(B)** Representative traces of SBI-553 and NTS perfusion **(C)** Neurotensin increases the sIPSC frequency in both males and females. This effect returns to baseline levels during the washout. Neurotensin induces a sex x drug interaction on **(D)** IPSC amplitude. **(E)** IPSC area in males and females however is not significantly different. **(F)** SBI-553 alone does not alter the IPSC frequency. SBI-553 is unable to block NTS-induced increases in IPSC frequency, but blocks NTS-induced increases in **(G)** IPSC amplitude and reduces the trend in **(H)** IPSC area in the males. Representative traces of **(I)** NTS perfusion in males and females. *p<0.05, **p<0.01

Since SBI-553 blocks NTS-mediated increases in GABAergic neurotransmission in the CeA_L_, we hypothesized that we would observe similar effects in ovBNST neurons. Like the CeA_L_, SBI-553 alone does not alter sIPSC frequency, amplitude, or area in either males or females. Unlike the CeA_L_, however, pre-treatment with SBI-553 did not block NTS-induced increases in sIPSC frequency (Fig. 3F; two-way repeated measures ANOVA; males: n= 8 cells, 5 mice; females: n= 5 cells, 4 mice; significant main effect of drug *F*_(1.245,13.70)_=17.51, *p*=0.0006, η^2^_p_= 0.614, ω^2^_p_= 0.717; no main effect of sex *F*_(1,11)_=0.334, *p*=0.575, and no drug x sex interaction *F*_(2,22)_=0.254, *p*=0.778; Tukey’s multiple comparisons test; baseline vs. SBI-553+NTS, *p*=0.0012, SBI-553 vs. SBI-553+NTS, *p*=0.0022). The cumulative distributions complement the sIPSC averages and show leftwards shifts in the interevent interval with SBI-553 and NTS (Sup Fig. 1K). Notably, SBI-553 prevents the NTS-induced sex x treatment interaction on sIPSC amplitude, (Fig. 3G; two-way repeated measures ANOVA; no main effect of drug *F*_(1.640,18.04)_=0.128, *p*=0.841; no main effect of sex *F*_(1,11)_=3.293, *p*=0.097, and no drug x sex interaction *F*_(2,22)_=0.169, *p*=0.845). sIPSC area also showed no effects (Fig. 3H; two-way repeated measures ANOVA; no main effect of drug *F*_(1.571,17.28)_=0.092, *p*=0.870; no main effect of sex *F*_(1,11)_=0.471, *p*=0.507, and no drug x sex interaction *F*_(2,22)_=0.359, *p*=0.702). These findings suggest that SBI-553 exerts region-specific effects on NTS modulation of GABAergic neurotransmission and that the sex-specific effects of NTS are due to divergent downstream signaling pathways in the BNST.

To further investigate the locus and mechanism of action of endogenous NTS signaling and the impact of biased signaling at NTSR1 within the CeA and BNST, we compared the protein expression of the early immediate gene cFos in tissue perfused 90 minutes after a single injection of SBI-553 (12 mg/kg) or vehicle (5% hydroxypropyl-β-Cyclodextrin) in behaviorally naïve male and female mice. SBI-553 causes a trend for an increase in cFos immunoreactivity in the CeA_C_ (Fig. 4A; two-way ANOVA; trending main effect of SBI-553 *F*_(1,25)_=3.588, *p*=0.070; no main effect of sex *F*_(1,25)_=0.323, *p*=0.575, and no SBI-553 x sex interaction *F*_(1,25)_=0.0009, *p*=0.977), but no significant changes in the CeA_M_ (Fig. 4B; two-way ANOVA; no main effect of SBI-553 *F*_(1,25)_=0.020, *p*=0.888; no main effect of sex *F*_(1,25)_=0.285, *p*=0.599, and no SBI-553 x sex interaction *F*_(1,25)_=0.715, *p*=0.406). Notably, SBI-553 significantly increases cFos immunoreactivity in the CeA_L_, an effect driven by the females (Fig. 4C; two-way ANOVA; significant main effect of SBI-553 *F*_(1,25)_=4.819, *p*=0.038 η^2^_p_= 0.161, ω^2^_p_= 0.116; no main effect of sex *F*_(1,25)_=0.368, *p*=0.550, and no SBI-553 x sex interaction *F*_(1,25)_=1.847, *p*=0.186; Bonferroni post-hoc test; males vehicle vs. SBI-553, *p*>0.999, females vehicle vs. SBI-553, *p*=0.042). These findings suggest that SBI-553 increases the activity of neurons within the CeA_L_, consistent with our electrophysiology data that SBI-553 reduces GABAergic input onto CeA_L_ neurons. Further, it suggests that NTS tone is significantly upregulated in the CeA_L_ of naïve females compared to naïve males. SBI-553 does not alter the cFos immunoreactivity of neurons in the dorsal BNST (Fig. 4E; two-way ANOVA; no main effect of SBI-553 *F*_(1,26)_=0.579, *p*=0.454; no main effect of sex *F*_(1,26)_=0.569, *p*=0.458, and no SBI-553 x sex interaction *F*_(1,26)_=0.474, *p*=0.497), ventral BNST (Fig. 4F; two-way ANOVA; no main effect of SBI-553 *F*_(1,26)_=0.239, *p*=0.629; no main effect of sex *F*_(1,26)_=1.173, *p*=0.289, and no SBI-553 x sex interaction *F*_(1,26)_=0.861, *p*=0.362), or the ovBNST (Fig. 4G; two-way ANOVA; no main effect of SBI-553 *F*_(1,26)_=0.0085, *p*=0.927; no main effect of sex *F*_(1,26)_=0.671, *p*=0.420; and no SBI-553 x sex interaction *F*_(1,26)_=0.3298, *p*=0.571), consistent with the electrophysiology data that SBI-553 only partially impacts on NTS-mediated changes to GABAergic neurotransmission in the ovBNST.

**Figure 4.**
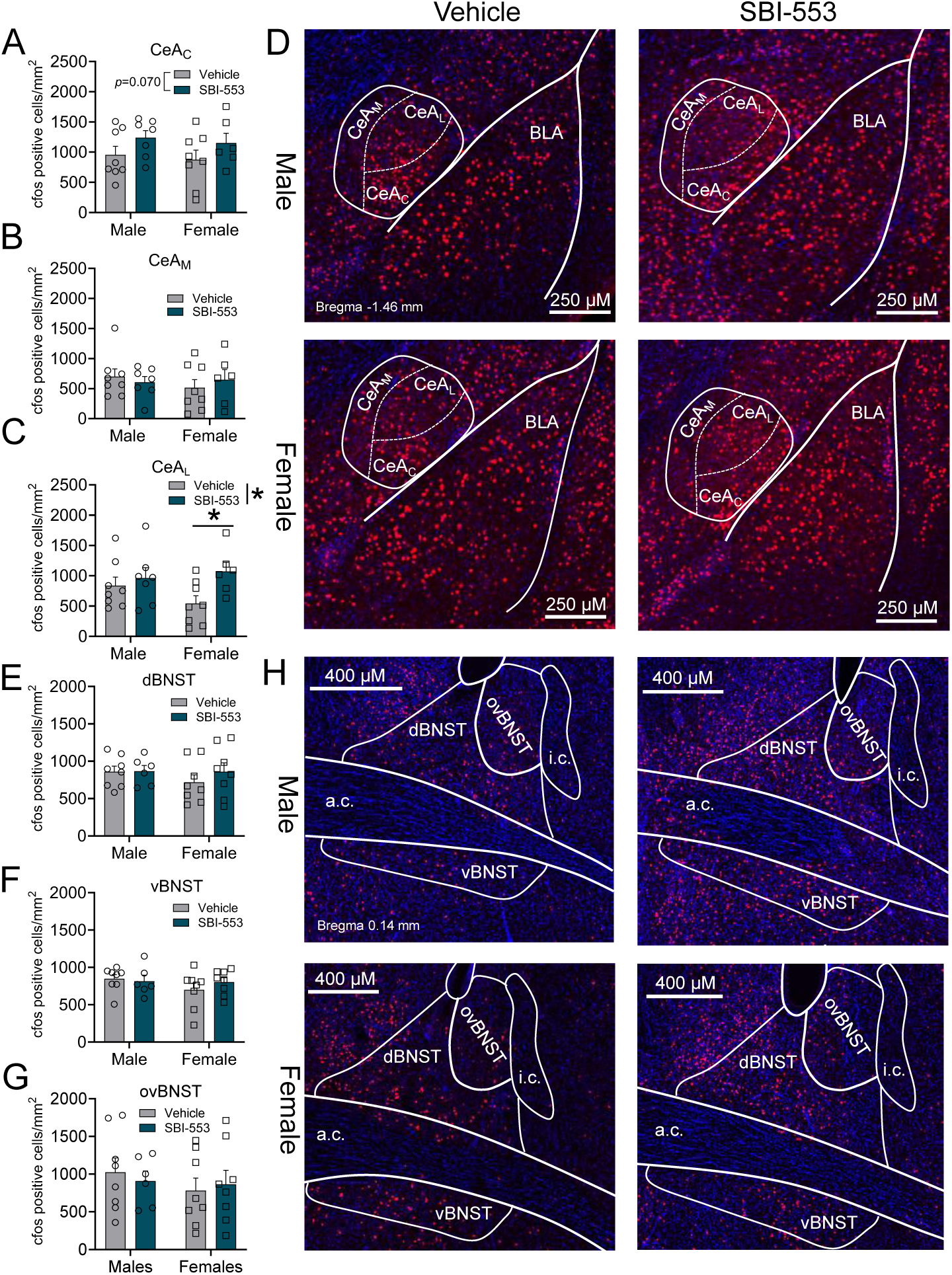
Single injection of SBI-553 alters the cFos immunofluorescence in the CeA, but not the BNST of naïve animals. **(A)** SBI-553 does not alter cFos expression in the CeC **(B)** or CeM of naïve animals **(C)** but selectively increases activity in the CeL of female mice compared to vehicle treatment. **(D)** Representative immunofluorescence images of cFos expression (red) and DAPI (blue) in the CeA. **(E)** SBI-553 has no impact on cFos activity in the dBNST, **(F)** vBNST, **(G)** or ovBNST of naïve male or female mice. **(H)** Representative immunofluorescence images of cFos expression in the BNST. Error bars represent average value ±SEM. *p<0.05

Next, we sought to probe the effect of systemic SBI-553 on feeding behavior and subsequent alterations to cFos expression in the CeA and BNST. Previously, our group found that SBI-553 (12 mg/kg, I.P.) reduces the consumption of highly palatable foods (FL) during the homecage re-feed post-test period in the NSFT [66] without affecting the latency to feed [66] suggesting that SBI-553 had effects on motivation to feed. Because the homecage re-feed is a control for the NSFT, we wanted to explore whether SBI-553 alters the motivational drive to eat without the confounding stress induced by the novel aversive environment. Ergo, we conducted the motivated homecage feeding assay (MHFA; see methods and Fig. 5A) to investigate how SBI-553 alters the consumption of a sugary, palatable food in fasted mice without prior exposure to a novel environment. Similar to our prior study, SBI-553 (12 mg/kg, I.P.) significantly reduces consumption of FLs in a 10-minute free-feeding test in both male and female mice (Fig. 5B; two-way ANOVA; significant main effect of SBI-553 *F*_(1,35)_=31.90, *p*<0.0001, η^2^ = 0.477, ω^2^ = 0.442; significant main effect of sex *F* =32.32, *p*<0.0001, η^2^ = 0.480, ω^2^ = 0.445; no SBI-553 x sex interaction *F*_(1,35)_=1.451, *p*=0.236; Bonferroni post-hoc test; males vehicle vs. SBI-553, *p*=0.005, females vehicle vs. SBI-553, *p*<0.0001,) compared to the vehicle control. Although females consumed more FL than males as a proportion of their bodyweight, this was due to female mice being smaller than males (Sup Fig. 2A; two-way ANOVA; no main effect of drug treatment group *F*_(1,35)_=1.567, *p*=0.2189; main effect of sex *F*_(1,35)_=71.51, *p*<0.0001, η^2^ = 0.671, ω^2^ = 0.709; no SBI-553 x sex interaction *F* =0.0048, *p*=0.9451). Moreover, when comparing grams of FL consumed during the 10-minute access period, males and females consumed similar amounts during the task under both conditions (Sup Fig. 2B; two-way ANOVA; significant main effect of SBI-553 *F*_(1,35)_=33.260, *p*<0.0001, η^2^ = 0.487, ω^2^ = 0.527; no main effect of sex *F* =2.610, *p*=0.115; no SBI-553 x sex interaction *F*_(1,35)_=0.0534, *p*=0.819).

**Figure 5.**
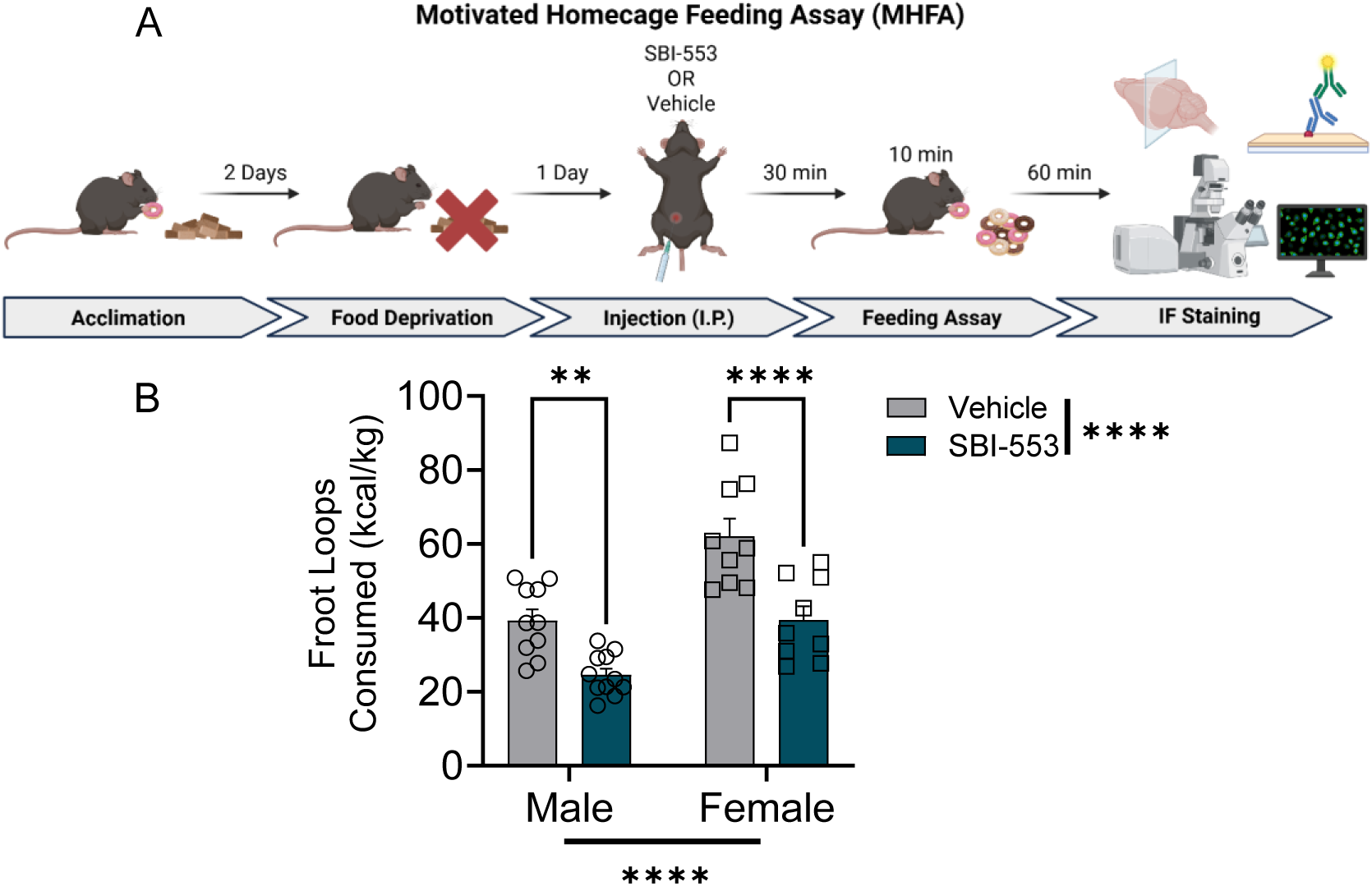
SBI-553 decreases palatable food consumption in both male and female mice in the Motivated Homecage Feeding Assay (MHFA). **(A)** Schematic of the MHFA timeline. **(B)** SBI-553 reduces palatable motivated feeding in both male and female mice. Error bars represent average value SEM. ** p<0.01, **** *p*<0.0001

To further examine potential mechanisms underlying SBI-553-induced changes in the MHFA, we quantified cFos expression in the CeA and BNST. In the MHFA, we found SBI-553 significantly increases the cFos expression in the CeA_C_ (Fig. 6A; two-way ANOVA; significant main effect of treatment *F*_(1,18)_=4.579, *p*=0.046, η^2^ = 0.203, ω^2^ = 0.140; no main effect of sex *F* =2.441, *p=*0.136; no sex x treatment interaction *F*_(1,18)_=2.591, *p*=0.125). Furthermore, females expressed lower cFos expression than males after vehicle treatment (Fig. 6A; Bonferroni post-hoc test; vehicle males vs females *p=*0.039). Conversely, no differences in expression were detected in the CeA_M_ (Fig. 6B; two-way ANOVA; no main effect of treatment *F*_(1,18)_=0.1617, *p*=0.692; no main effect of sex *F*_(1,18)_=0.3260, *p*=0.575; no sex x treatment interaction *F*_(1,18)_=2.713, *p*=0.117) or the CeA_L_ (Fig. 6C; two-way ANOVA; no main effect of treatment *F*_(1,18)_=0.6517, *p=*0.430; no main effect of sex *F*_(1,18)_=0.7824, *p*=0.388; no sex x treatment interaction *F*_(1,18)_=0.3168, *p*=0.581). SBI-553 also does not alter cFos expression in the dorsal (Fig. 6E; two-way ANOVA; no main effect of treatment *F*_(1,17)_=0.4362, *p*=0.518; no main effect of sex *F*_(1,17)_=0.368, *p*=0.522; no sex x treatment interaction *F*_(1,17)_=0.0188, *p*=0.893), ventral (Fig. 6F; two-way ANOVA; no main effect of treatment *F*_(1,17)_=0.261, *p*=0.616; no main effect of sex *F*_(1,17)_=0.0928, *p*=0.764; no sex x treatment interaction *F*_(1,17)_=0.419, p=0.526), or oval (Fig. 6G; two-way ANOVA; no main effect of treatment *F*_(1,17)_=0.0676, *p*=0.780; no main effect of sex *F*_(1,17)_=0.0185, *p*=0.904; no sex x treatment interaction *F*_(1,17)_=0.0169, *p*=0.898) BNST subregions. These findings align with the cFos data in naïve animals, where SBI-553 produces a trend for an increase in cFos in the CeA_C_ and no effect in the BNST.

**Figure 6.**
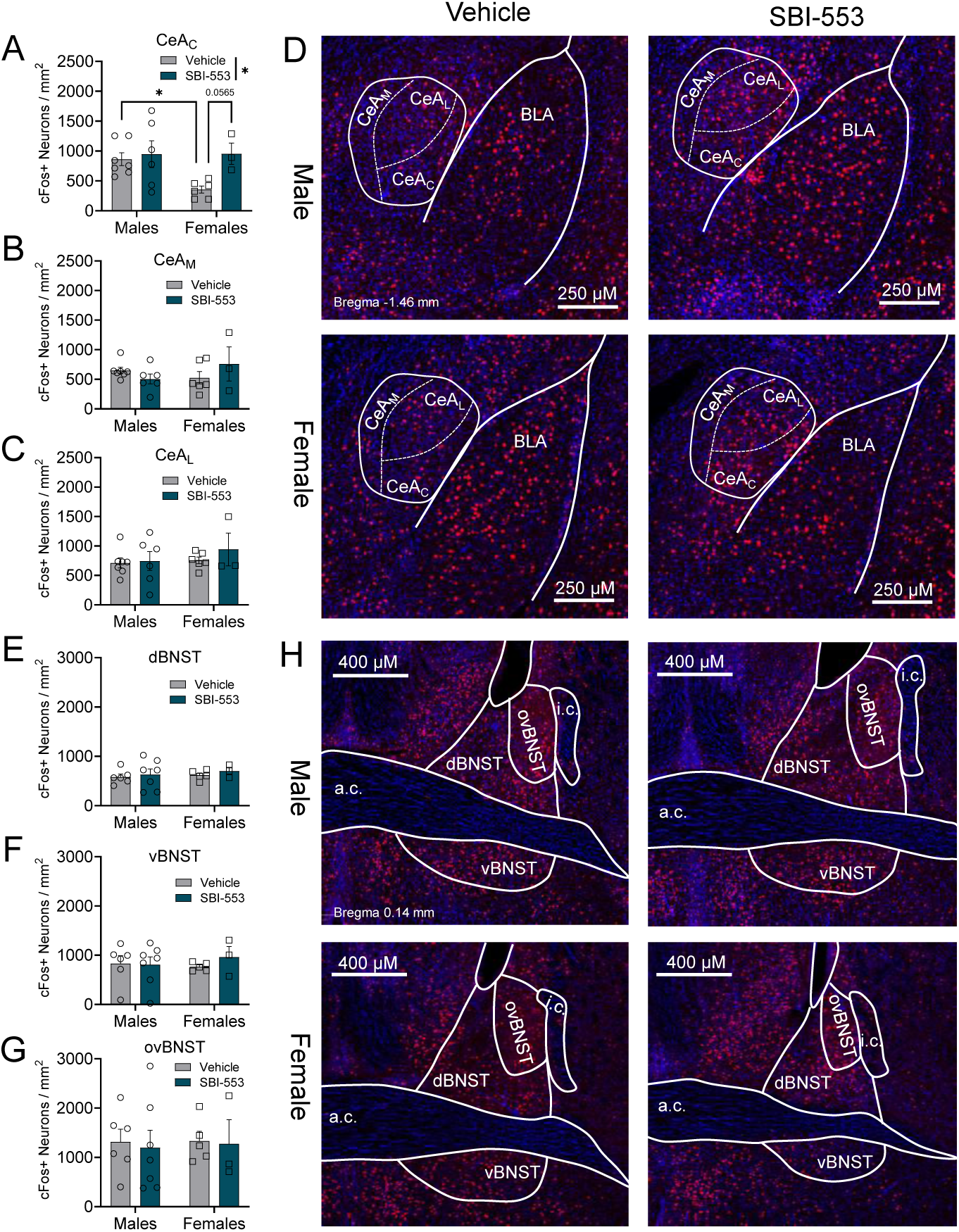
SBI-553 alters the cFos immunoflourescence in the CeA, but not the BNST after the MHFA. **(A)** SBI-553 significantly increases cFos expression in the CeAC, especially in females, **(B)** but not the CeM **(C)** or CeAL of any mice after the MHFA compared to vehicle treatment. **(D)** Representative immunofluorescence images of cFos expression (red) and DAPI (blue) in the CeA. **(E)** SBI-553 has no impact on cFos activity in the dBNST, **(F)** vBNST, **(G)** or ovBNST of male or female mice after the MHFA. **(H)** Representative immunofluorescence images of cFos expression in the BNST. Error bars represent average value SEM. * *p*<0.05

In addition to the extended amygdalar regions already described, we also performed a preliminary investigation the impact of SBI-553 signaling in two adjacent brain regions with established roles in NTS signaling: the lateral hypothalamus (LH) and medial/lateral preoptic areas (mPOA/lPOA). The LHA contains a large population of NTS-expressing neurons that integrate metabolic signals to regulate feeding, energy expenditure, and mesolimbic dopamine signaling [79,80], while the preoptic area expresses both NTS and NTSR1 and mediates NTS-dependent regulation of thermoregulation and energy expenditure [81,82]. As such, we wanted to investigate whether SBI-553 altered activity beyond our primary extended amygdalar regions of interest. In the LH, after the MHFA task, we observed a significant interaction between sex and treatment (Fig. S4A; two-way ANOVA; sex x treatment interaction F(1,16)=7.535, p=0.014, η^2^_p_= 0.320, ω^2^_p_= 0.246; trending main effect of treatment F(1,16)=4.004, p=0.063; no main effect of sex F(1,16)=0.3738, p=0.550). Post-hoc analysis revealed that SBI-553 significantly increased cFos expression in females (Bonferroni post-hoc test; vehicle vs. SBI-553 p=0.008) but not males (vehicle vs. SBI-553 p=0.557), and that females exhibited significantly higher cFos expression than males following SBI-553 treatment (males vs. females p=0.042) with no sex difference under vehicle conditions (males vs. females p=0.121). Conversely, in the preoptic subregions, SBI-553 did not alter cFos expression in the mPOA (Fig. S4C; two-way ANOVA; no main effect of treatment F(1,17)=2.608, p=0.125; no main effect of sex F(1,17)=1.687, p=0.211; no sex x treatment interaction F(1,17)=1.327, p=0.265) or the lPOA (Fig. S4D; two-way ANOVA; no main effect of treatment F(1,17)=0.4394, p=0.516; no main effect of sex F(1,17)=0.009285, p=0.924; no sex x treatment interaction F(1,17)=1.218, p=0.285). These data suggest that in fasted mice systemic SBI-553 engages the LH in a sex-dependent manner, with females showing significantly elevated cFos activity following SBI-553 treatment, but does not broadly alter activity across adjacent preoptic regions.

Since animals in both the NSFT and MHFA are food-restricted for 20-24 hours prior to testing, we examined the impact of food availability on SBI-553 induced changes in FL consumption using the FvFHFA. The FvFHFA allows us to explore the role of energy balance on feeding behavior and assess whether a caloric deficit shifts the motivational state and influences SBI-553-induced reductions in FL consumption. In a 3-way RM-ANOVA, we did not observe a main effect of sex (p=0.392) in feeding behavior in this assay, so we collapsed sex to increase power for analysis. In this experiment, all animals were treated for one week with vehicle, and the other week with SBI-553, counterbalanced, allowing for a within-animal comparison of SBI-553 treatment on FL consumption. We did not observe an effect of treatment sequence (Fig 7D; two-way RM ANOVA; significant main effect of SBI-553 *F*_(1,24)_=21.25, *p*=0.0001, η^2^_p_= 0.470, ω^2^_p_= 0.438; no main effect of treatment order *F*_(1,24)_=1.459, *p*=0.239; no treatment x treatment order effect *F*_(1,24)_=0.0398, *p*=0.844). As expected, fasted animals consumed significantly more FLs during the 10-minute period when given vehicle than fed animals (Fig. 7B; two-way RM ANOVA; significant main effect of feeding condition *F*_(1,24)_=10.89, *p*=0.003, η^2^_p_= 0.512, ω^2^_p_= 0.481, significant main effect of SBI-553 *F*_(1,24)_=25.560, p<0.0001, η^2^_p_= 0.516, ω^2^_p_= 0.486; and a significant feeding condition x SBI-553 interaction F_(1,24)_=4.550, p=0.0433, η^2^_p_= 0.160, ω^2^_p_= 0.120; Bonferroni post-hoc test; vehicle fed vs fasted *p*=0.0005). SBI-553 significantly attenuates FL consumption in fasted animals, normalizing their FL intake to that of fed animals (Fig. 7B; Bonferroni post-hoc test; vehicle fasted vs. SBI-553 fasted p<0.0001, SBI-553 fed vs fasted p=0.240). Because there was a feeding status x treatment interaction, and no effect of treatment order (*p*=0.240), we calculated the within-animal difference (Δ kcal/kg consumption) after treatment with SBI-553–vehicle. The reduction in consumption induced by SBI-553 was larger in the fasted group vs the fed group (Fig. 7C; unpaired t test; fed vs fasted t_(24.00)_=2.133, *p*=0.043, Cohen’s d =0.836), suggesting that SBI-553 biased modulation of NTSR1 is stronger when metabolic or motivational state drives feeding behavior.

**Figure 7.**
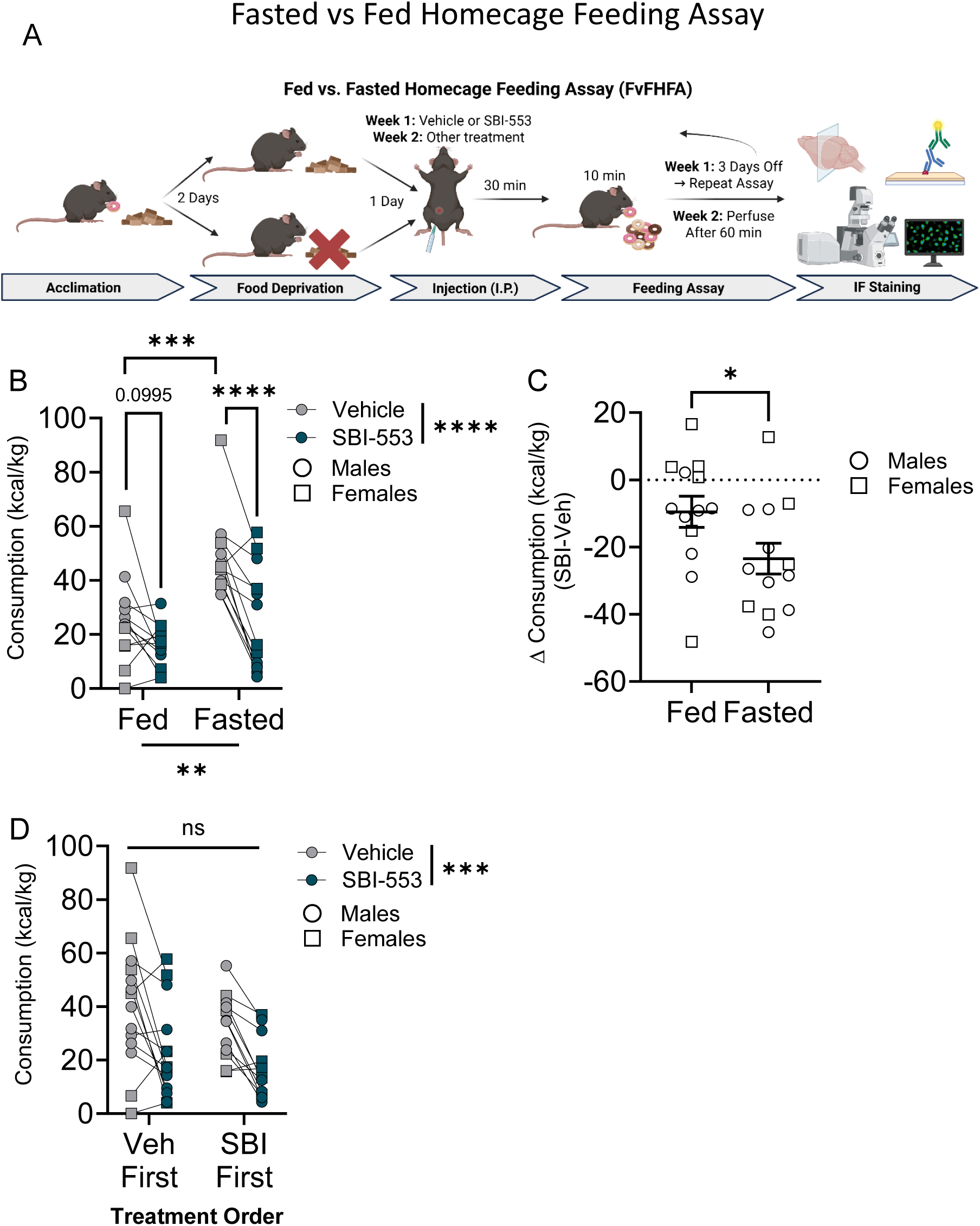
SBI-553 reduces palatable FL consumption in both fed and fasted conditions. **(A)** Schematic of the FvFHFA timeline. **(B)** SBI-553 significantly reduces feeding behavior in both fed and fasted feeding states, with **(C)** a larger reduction in in the fasted animals. **(D)** There was no difference in treatment order on reductions of consumption across feeding conditions. Error bars represent average value ± SEM. * *p*<0.05, ** *p*<0.01, *** *p*<0.001, **** *p*<0.0001

In the FvFHFA, we found no main effects when comparing all 3 conditions (sex, treatment, and feeding condition) (Sup Fig. 3). Therefore, we collapsed sex to increase the statistical power to investigate the differences in cFos expression between treatment and feeding conditions across amygdalar subregions. In the FvFHFA, SBI-553 significantly increases cFos expression in both the CeA_C_ (Fig. 8A; two-way ANOVA; significant main effect of treatment *F*_(1,19)_=4.571, *p*=0.046, η^2^_p_= 0.194, ω^2^_p_= 0.134; no main effect of feeding condition *F*_(1,19)_=0.1352, *p*=0.717; no feeding condition x treatment interaction *F*_(1,19)_=0.3461, *p*=0.563) and CeA_L_ (Fig. 8C; two-way ANOVA; significant main effect of treatment *F*_(1,19)_=5.031, *p*=0.037, η^2^_p_= 0.209, ω^2^_p_= 0.149; no main effect of feeding condition *F*_(1,19)_=2.236, *p*=0.151; no feeding condition x treatment interaction *F*_(1,19)_=0.1153, p=0.738), without altering expression in the CeA_M_ (Fig. 8B; two-way ANOVA; no main effect of treatment *F*_(1,19)_=3.283, *p*=0.086; no main effect of feeding condition *F*_(1,19)_=1.850, *p*=0.190; no feeding condition x treatment interaction *F*_(1,19)_=0.0690, *p*=0.796). SBI-553 does not impact cFos expression in the dBNST (Fig. 8E; two-way ANOVA; no main effect of treatment *F*_(1,13)_=0.1222, *p*=0.732; no main effect of feeding condition *F*_(1,13)_=0.2008, *p*=0.662; no feeding condition x treatment interaction *F*_(1,13)_=0.9475, *p*=0.348). In the vBNST, fasted animals have higher cFos expression than their fed counterparts after SBI-553 treatment (Fig. 8F; two-way ANOVA; significant main effect of feeding condition *F*_(1,14)_=5.762, *p*=0.031, η^2^_p_= 0.292, ω^2^_p_= 0.209; no main effect of treatment *F*_(1,14)_=0.774, *p*=0.394; no feeding condition x treatment interaction *F*_(1,14)_=3.321, *p*=0.090; Bonferroni post-hoc test; SBI-553 fed vs fasted *p*=0.035). Additionally, an SBI-553 x feeding condition interaction was observed observed in the ovBNST, with a trend for higher cFos expression in fed animals receiving SBI-553 compared to vehicle and no difference in the fasted animals (Fig. 8G; two-way ANOVA; no main effect of treatment *F*_(1,14)_=1.088, *p*=0.315; no main effect of feeding condition *F*_(1,14)_=0.104, *p*=0.752; significant feeding condition x treatment interaction *F*_(1,14)_=4.912, p=0.044, η^2^_p_= 0.260, ω^2^_p_= 0.179; Bonferroni post-hoc test; trend in Fed vehicle vs SBI-553 *p*=0.055). Together, these results indicate that subregions are differentially sensitive to SBI-553 administration depending on whether the animal is fed or fasted.

**Figure 8.**
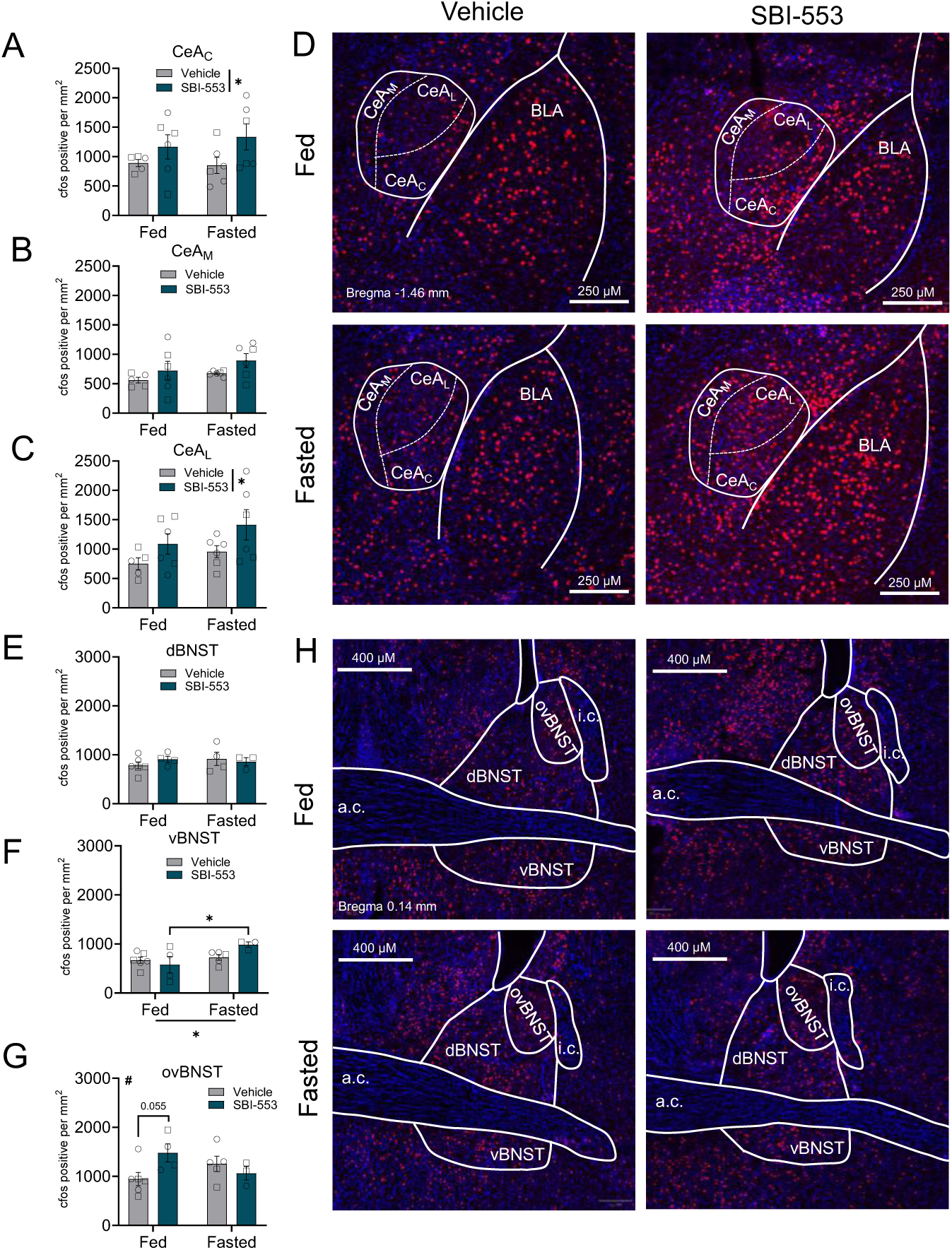
SBI-553 alters the cFos immunoflourescence in the CeA and the ovBNST after the FvFHFA. **(A)** SBI-553 significantly increases cFos expression in the CeAC and **(C)** CeAL, **(B)** but not the CeAM compared to vehicle treatment across feeding conditions. **(D)** Representative immunofluorescence images of cFos expression (red) and DAPI (blue) in the CeA. **(E)** SBI-553 has no impact on cFos activity in the dBNST in either fed or fasted animals in the FvFHFA. **(F)** In the vBNST, fasted animals showed higher levels of cFos activity than fed animals, which was significantly different between animals receiving vehicle or SBI-553. **(G)** ovBNST cFos activity not impacted by SBI-553 treatment in fasted animals but showed trending elevations in animals receiving SBI-553 in a fed state compared to vehicle, with a significant interaction between SBI-553 and feeding condition (two-way ANOVA, interaction p=0.04). **(H)** Representative immunofluorescence images of cFos expression in the BNST. Error bars represent average value ± SEM. * *p*<0.05, # interaction *p*<0.05

Interestingly, when looking at cFos activity in the LH after FvFHFA, no differences in expression were detected following the FvFHFA (Fig. S5A; two-way ANOVA; no main effect of treatment F(1,15)=0.5537, p=0.468; no main effect of sex F(1,15)=0.02352, p=0.880; no sex x treatment interaction F(1,15)=0.7116, p=0.412). However, in the mPOA, SBI-553 significantly increased cFos expression (Fig. S5C; two-way ANOVA; significant main effect of treatment F(1,10)=7.801, p=0.019, η^2^ = 0.438, ω^2^ = 0.327; no main effect of sex F(1,10)=0.1920, p=0.671; no sex x treatment interaction F(1,10)=0.4044, p=0.539). No differences were detected in the lPOA (Fig. S5D; two-way ANOVA; no main effect of treatment F(1,10)=1.878, p=0.201; no main effect of sex F(1,10)=0.7041, p=0.421; no sex x treatment interaction F(1,10)=0.0001, p=0.992).

## Discussion

### Summary of Results

GPCRs are the largest and most diverse family of membrane-bound receptors, and drugs that modulate GPCR signaling constitute a significant portion of currently available pharmacotherapies across a range of disease states [83]. Despite their clinical success, the lack of receptor subtype selectivity and off-target effects are disadvantages of traditional GPCR agonists and antagonists that commonly produce unwanted side effects. Biased allosteric modulators (BAMs), compounds that target sites on receptors outside the orthosteric pocket to alter intracellular signaling pathways, have recently emerged as novel tools that modulate GPCR signaling to produce therapeutic benefit while mitigating undesired side effects [84]. One such ligand is SBI-553, an NTSR1 b-arrestin BAM, which has been shown to attenuate cocaine self-administration and alcohol consumption in mice without the side effects associated with NTSR1 agonists [33,55]. Although preclinical data highlight the therapeutic potential of BAMs, there has been limited investigation into the specific cellular and neurophysiological effects of NTSR1 BAMs in brain regions governing motivated behaviors [84]. Here, we investigated the impact of NTS and the NTSR1 BAM, SBI-553, on *ex vivo* GABAergic signaling and cellular activation in naïve mice across the extended amygdalar subregions of the CeA and BNST. Further, we probed whether SBI-553 alters the consumption of sugary, palatable foods and cFos activity in these regions across feeding states.

### NTS and SBI-553 differentially alter GABAergic signaling across Central Amygdala subregions in naïve mice

As prior work has found the effects of NTS to robustly modulate inhibitory neurotransmission in the extended amygdala [71,73,74], we focused on IPSCs in our current study. Using whole-cell voltage clamp electrophysiology to record sIPSCs in naïve male and female mice, we show that CeA and BNST subregions are differentially responsive to NTS and SBI-553 application. Specifically, we demonstrate that exogenous NTS enhances GABAergic signaling in the CeA_L_, but not in the CeA_M_. This subregion-specific effect is consistent with the known distribution of NTS and NTSR1 within the CeA, where NTS and NTSR1 mRNA are most densely localized in the CeA_L_ [78–80]. Multiple groups have reported high colocalization of Sst, Crf, and Tac2 in CeA_L_^NTS^, with little colocalization detected in CeA_M_^NTS^ [19,86], suggesting that CeA^NTS^ have distinct molecular profiles across CeA_M_ and CeA_L_ subregions. The CeA also contains significant variability in the electrophysiological properties of neurons within its subregions [85]. NTSR1 activation excites CeA_L_ neurons via a G_q_-mediated and PLC-dependent mechanism that inhibits inwardly rectifying potassium channels. Notably, NTS-induced excitation of CeA_L_ neurons has been shown to enhance GABAergic transmission (sIPSCs) onto CeA_M_ neurons [85]. In contrast, our findings show that NTS does not alter GABAergic signaling in the CeA_M_, suggesting that the CeA_M_ is a downstream target of NTS-mediated CeA_L_ inhibition rather than a direct site of NTS action. The discrepancy between these studies could be explained by several factors: (1) We recorded from adult C57BL6/J mice, whereas Lei and Hu recorded from adolescent 3-4 week old Sprague-Dawley rats. The NTS system is differentially expressed across species and throughout development [87–90], and these developmental differences may, in part, be regulated by DNA methylation and other signaling motifs in the *Nts* promoter region. (2) We applied NTS via bath application versus focal application with a picospritzer, which may differentially engage local microcircuits and afferent projections. Although our findings suggest that NTS has no effect on GABAergic signaling in the CeA_M_, females have a higher baseline sIPSC frequency than males which could be developmentally regulated. Similar sex differences in basal GABAergic transmission within the CeA_M_ have previously been reported [75]. The Roberto lab has previously shown that sex differences in CeA physiology vary by subregion and modulator: CeA_L_ neurons in female rats showed blunted corticosterone sensitivity, whereas CeA_M_ responses were unchanged, suggesting that circuit properties may be impacted by sex in a subregion-dependent manner [23]. We have also previously demonstrated that chronic changes in GABAergic tone can reshape postsynaptic responses in a sexually divergent manner, where reductions in GABAergic neurotransmission induced by vGAT knockdown in CeA^NTS^ neurons increase the sIPSC decay exclusively in males [34]. Together, these findings complement our current results and suggest that baseline GABAergic signaling may reflect sex differences in the density of GABAergic afferents to CeA_M_, GABA release probability, or GABA_A_ subunit composition in the CeA_M_.

We find that preincubation with SBI-553 prevents the NTS-induced increase in GABA transmission in the CeA_L_, likely via enhanced presynaptic release. These findings are consistent with previous reports suggesting that NTS activates NTSR1 and downstream G_q_-coupled signaling pathways in the CeA_L_ to alter GABA release [85]. We have previously demonstrated minimal overlap between NTS and NTSR1 expression in CeA neurons [40], suggesting that NTSR1 modulates presynaptic GABA input from local interneurons or afferent projections. These results support findings from the Hnasko lab, where SBI-553 prevented dopamine neuron firing and terminal release in the nucleus accumbens using *ex* and *in vivo* preparations [84].

### SBI-553 does not inhibit NTS-induced increases in GABAergic signaling in the BNST

We previously observed a NTSR1-mediated sustained increase in electrically and optically evoked IPSCs in the ovBNST following NTS application, indicative of inhibitory long-term potentiation (iLTP) of CeA-BNST inputs. Specifically, we found the nonselective NTSR antagonist SR142948A, but not the NTSR2-selective antagonist NTRC844 blocked the NTS-mediated enhancement of GABA signaling. In contrast to the increase we observed in evoked and now here in sIPSCs with NTS, the NTSR2-selective agonist levocabastine reduced the IPSC amplitude in ∼40% of the cells assayed [40]. This suggests that the overall effect of endogenously released NTS or a 1 µM bath application is an enhancement of GABAergic signaling, although this balance could be modified under behavioral challenges like stress, drug exposure, or metabolic disease. In the present study, we found that NTS-mediated enhancement of sIPSC frequency is transient, suggesting that other neuropeptides and/or signaling molecules released during evoked transmission or cellular depolarization are necessary to sustain the NTS-mediated iLTP. Additionally, the recordings in our previous study were also conducted on a reverse light cycle, and data from the Kash lab has demonstrated that BNST neuropeptide Y modulation alters inhibitory neurotransmission differentially based on the cells being recorded from in the light or dark cycle [91], which may suggest circadian adaptation in this signaling.

Contrary to the CeA_L_, we found that surprisingly SBI-553 pretreatment is unable to block NTS-mediated increases in sIPSC frequency in the ovBNST of male and female mice. Although the CeA_L_ and ovBNST are generally homologous structures with similar neuropeptide and signaling molecule expression [42,80,89], these findings highlight differential responsivity to modulation of the neurotensinergic system across regions of the extended amygdala. Additionally, NTS does not change the sIPSC amplitude in either sex in the CeA_L_ but induced a sex x drug treatment interaction the sIPSC amplitude in the ovBNST, such that neurotensin significantly increased the amplitude of the sIPSC in males. These findings suggest that NTSR1 activation may promote greater GABAergic inhibition in the ovBNST in males than females, possibly promoting sex differences across a range of behaviors. Notably, while SBI-553 did not prevent NTS-mediated increases in sIPSC frequency, it does block the sex x drug treatment interaction in sIPSC amplitude in male mice. These data suggest that NTSR1 signals via at least two distinct mechanisms in the ovBNST. Additionally, it was surprising the sIPSC area did not reflect the same changes as frequency or amplitude. This may, however, indicate that the effects on frequency and amplitude may be occurring on different synapses. Our findings illustrate that future studies in both sexes are necessary to examine the effects of NTS and SBI-553 on specific cell types, incorporating both local BNST circuits and GABAergic afferant projections to the ovBNST, and at different circadian time points in the BNST during different feeding states/metabolic challenges. Additionally, our future studies plan to explore how NTS and SBI-553 alter cell firing and membrane voltage (current clamp), as well as how SBI-553 application subsequent to NTS affects cellular potential and synaptic transmission.

### SBI-553 alters the activation of CeA_L_ neurons of naïve mice in vivo

Based on our *ex vivo* findings of NTS and SBI-553-mediated changes in GABAergic neurotransmission in the CeA and BNST, we sought to investigate how activity in extended amygdala subregions was impacted *in vivo* using cFos immunofluorescence staining as an index of neuronal activity. In naïve mice, we found that SBI-553 significantly increases cFos expression in the CeA_L_, an effect driven by the female mice. These differences imply possible sex differences in either SBI-553 signaling, G-protein coupling, endogenous NTS tone, and/or afferent inputs to the CeA_L_ in male and female mice. Indeed, the neurotensin system is well known to be regulated by sex [93], and our previous study found SBI-553 altered behavioral and physiological responses to alcohol in a sex-dependent manner [66]. Although SBI-553 antagonizes G_q_-mediated signaling and promotes the b-arrestin pathway [56], SBI-553 also activates Gα_15_, multiple G_i/o_ subtypes, G_12/13_, and b-arrestin in the absence of NTS; however, these effects are modulated when NTS is present, and in particular G_12/13_ signaling is enhanced by SBI-553 [60], and G_12/13_ signaling is implicated in metabolic disease [94]. This may suggest that distinct intracellular signaling cascades could result in increased cFos expression in the CeA_L_ [57]. Additionally, these results may reflect sex differences in basal NTS tone, aligning with previous work [93,95]. Thus, the larger SBI-553-induced increases in CeA_L_ cFos expression in females may arise from baseline sex differences in endogenous NTS tone, NTSR1 expression, altered G-protein/b-arrestin signaling, or a combination thereof.

### SBI-553 reduces palatable food consumption in both food restricted and ad lib fed mice

The NTS system has been implicated in feeding and consummatory behavior for decades, with previous *in vivo* findings demonstrating that central and peripheral administration of NTS suppresses feeding behaviors in rats [7,96]. More recent studies have demonstrated that NTS is not exclusively an anorectic peptide, and that its impact may differ across a variety of feeding states, including fasted or obesity-induced motivated feeding behaviors [97–99]. It has also been shown in humans that circulating pro-NTS (the precursor to NTS) is elevated in insulin-resistant and clinically obese individuals and that high pro-NTS levels increase the likelihood of developing obesity [6,7]. Moreover, medical interventions for obesity (i.e., gastric bypass, gastric band, and caloric deficit) have differential impacts on restoring plasma NTS levels [100–102]. To further explore the role of NTS modulation on palatable food consumption, we investigated the effects of systemic SBI-553 on feeding behaviors and subsequent cFos activity. In addition to reducing binge alcohol consumption and preference, our prior research found that SBI-553 reduces post-test FL consumption in the NSFT in fasted mice [33]. This effect was notable because SBI-553 does not alter binge sucrose consumption (3% sucrose in water) [33]. However, these animals had *ad lib* access to food and water and were not exposed to an aversive, novel environment such as the NSFT. To address these confounds, we conducted the MHFA in fasted animals, which showed that SBI-553 reduced FL consumption compared with vehicle controls without prior exposure to a novel and stressful environment. The MHFA leverages 2 forms of motivation: (1) decreased energy balance and (2) highly palatable food. To test if an energy deficit was required for the action of SBI-553, we used the FvFHFA to compare the actions of SBI-553 in both food-restricted and *ad lib* fed animals using a within-animal design. Our data indicates that SBI-553 reduced feeding in both fed and fasted animals; however, the effects were significantly larger in mice with a caloric deficit and only trending in the fed mice. FLs are highly palatable, calorically dense, and rewarding food, and FL consumption engages hedonic feeding circuits [103]. The observation that SBI-553 reduces consumption across metabolic states suggests NTSR1 modulation may impact the hedonic process that drives consumption of palatable food independent of caloric need. There is evidence that IPAC^NTS^ neurons encode dietary preference for energy-dense foods and bidirectionally regulate hedonic feeding [98].

Furthermore, we have demonstrated that optogenetic stimulation of PBN-projecting CeA^NTS^ neurons promotes consumption of palatable (alcohol, sucrose, and saccharin), but not neutral (water) or aversive (quinine) fluids [13]. The larger effect of SBI-553 on FL consumption in food-deprived animals may therefore reflect a heightened motivational drive in the fasted animals because of their negative energy balance, which SBI-553 blunts more strongly than in fed animals. NTS signaling has been shown to play a critical role in energy balance regulation, particularly through NTS neurons in the LH that modulate feeding behavior and body weight [15,98]. Future studies examining how SBI-553 modulates interoceptive feeding signals that are disrupted in states of obesity will be critical for evaluating the therapeutic potential of NTSR1 allosteric modulation in metabolic diseases and motivated consummatory behaviors.

### SBI-553 increases cFos activity differently across CeA subregions and feeding state

Across our *in vivo* investigations, we observed increases in cFos activity in the CeA_L_ and CeA_C_ while CeA_M_ activity remained unchanged. This subregional pattern was consistent regardless of whether animals were behaviorally naïve, food-restricted (MHFA), or under different feeding states (FvFHFA), suggesting that SBI-553 produces stable activation signatures across lateral and capsular CeA subregions. In the BNST, however, differences in cFos expression emerged only when feeding conditions were directly compared in the FvFHFA, indicating that BNST responsivity to SBI-553 is dependent on the animal’s metabolic state. These findings align with recent work demonstrating that the ovBNST contains functionally opposing neuronal populations that are differentially modulated by metabolic state. Wang et al. [105] identified an ovBNST microcircuit in which PKC-δ⁺ neurons suppress feeding via monosynaptic GABAergic inhibition of LHA-projecting vlBNST neurons, whereas activation of those vlBNST neurons promotes food intake. In our FvFHFA data, SBI-553 increased cFos in the ovBNST of fed but not fasted animals, while vBNST cFos was elevated in fasted animals treated with SBI-553 relative to fed. This opposing pattern across BNST subregions is consistent with the inhibitory relationship between ovBNST and vlBNST. Recently, Kandil et al. identified a Vipr2⁺ population within the ovBNST which promotes feeding, is robustly activated by food restriction, projects to the parasubthalamic and paraventricular nuclei rather than LHA, and is largely non-overlapping with PKC-δ⁺ neurons [106]. The coexistence of these functionally opposing populations within the ovBNST suggests that the state-dependent cFos changes we observed may reflect differential recruitment of distinct cell types under different feeding conditions. Our cFos findings provide converging pharmacological evidence for this functional architecture, suggesting that NTSR1 modulation by SBI-553 interacts with the motivational state of the animal to shift the balance of activity across these BNST subregions. Interestingly, our preliminary exploration of the LHA and mPOA shows that SBI-553 increases cFos in the LHA exclusively in females, and there was a trend in the mPOA driven by females in the MHFA when all animals were fasted. Meanwhile, in the sex-collapsed FvFHFA, SBI-553 increases cFos in both the fed and fasted conditions in the mPOA with significant posthoc comparison between vehicle and SBI-553 in the fed condition. We do caution the reader that these preoptic/hypothalamic data are likely underpowered, and further investigation will likely yield additional insight. These data emphasize that future explorations of NTS/SBI-553 modulation of this interrelated circuitry should be done in intersectional genetic/projection manner.

### Mechanistic uncertainty and the complexity of systemic SBI-553, study limitations

This work adds to the growing body of literature suggesting that NTS modulation impacts aspects of food consumption, which may be related to hedonic features of the food and internal state of the animal. However, SBI-553 is neither a traditional agonist nor antagonist of NTSR1, and its signaling profile is complex. NTSR1 is canonically G_q_-coupled, however it can potently activate many other non-visual G_α_ proteins. [59]. Recent evidence from Moore et al. demonstrates that in addition to promoting beta-arrestin association, SBI-553 acts as a molecular ‘bumper’ and ‘glue’, inhibiting and promoting G protein binding in a subtype-specific manner [61]. While our study examined regional differences within the CeA and BNST, we did not examine effects on specific cell types, which may have differential expression of G-proteins and β-arrestin. Future studies will need to examine additional brain regions, focus on cell types within these brain regions, and additional microcircuitry elements. Although we examined sex as a biological variable in our studies here and reported sex differences in some assays, we note that some of our histology in the feeding assays is underpowered to detect sex differences and we caution misinterpreting that sex differences do not exist, in particular in the FvFHFA cFos data. Indeed, it is well appreciated that there are sex differences in both feeding behavior under different motivational drives [107–109], and within the NTS system which can be regulated by non-canonical estrogen signaling [95]. Additionally, although NTS and NTSR1 are expressed in many brain regions, including the CeA and BNST, the predominant source of NTS is in the intestine, and NTSR1 is expressed in other organs, including the pancreas [110]. Importantly, peripheral NTS also regulates food intake through both vagal and humoral routes [111], indicating that systemic SBI-553 may be modulating neurotensinergic signaling not only within the circuits examined here, but also at peripheral sites where gut-brain NTS communication is engaged. Disentangling the relative contributions of central versus peripheral NTSR1 modulation to the behavioral effects of SBI-553 will be an important goal for future studies.

### Conclusions and Future Directions

In summary, our findings demonstrate that NTS modulates GABAergic transmission in similar but divergent ways in the CeA_L_ and the ovBNST. Future studies will need to address cell type effects recording from both NTS and NTSR1 expressing cells. Additionally, future work should robustly probe the synaptic mechanism using miniature IPSC recordings, and electrically and optically evoked recordings. SBI-553 modulates GABAergic signaling in a subregion-specific manner within the extended amygdala, blocking NTS-induced increases in inhibitory tone in the CeA_L_ but has minimal impact on NTS-induced elevation of inhibitory transmission in the ovBNST. These data complement our *in vivo* findings, in which SBI-553 preferentially alters neuronal activation in the CeA_L_ and CeA_C_ but not the CeA_M_, further supporting the occurrence of functionally distinct neurotensinergic signaling across CeA subregions. In the BNST, SBI-553 alters neuronal activation in both the ovBNST and vBNST only when probing the feeding state, with no effects observed in naïve or fasted animals, suggesting that NTSR1 modulation in the BNST may be gated by the motivational context in which feeding occurs. Together, these data position the extended amygdala as a site where NTS signaling integrates interoceptive and motivational information to shape consummatory behavior. In the future, we will assess the effect of NTSR1 modulation on physiology in defined cells and circuits, considering hormonal status in male and female mice and circadian rhythms. Additional future studies employing different feeding paradigms, including operant consumption, will be valuable for disentangling the motivational versus consummatory components of SBI-553’s impacts on feeding. Additionally, intra-amygdalar infusions of SBI-553 will be an important future direction to dissect the relative contributions of central versus peripheral NTSR1 modulation to the behavioral effects observed here. Finally, examining SBI-553 in models of diet-induced obesity, in which interoceptive feeding signals and NTS tone are disrupted, will be critical for evaluating the translational potential of NTSR1 allosteric modulation for dysregulated feeding behaviors. In conclusion, we believe that biased modulation of NTSR1 may prove to be a useful pharmacotherapeutic for consummatory disorders.

## Acknowledgements

We thank Marc Louis for technical assistance with experiments.

This manuscript was supported by R01AA026363 (ZAM), T32AA007573 (SES), T32GM135095 (ARB), R01DA061773 (LMS), and R01DK103808 (GML).

Microscopy was performed at the UNC Hooker Imaging Core Facility, supported in part by P30 CA016086 Cancer Center Core Support Grant to the UNC Lineberger Comprehensive Cancer Center

## Author contributions

Conceptualization – SES, ARB, LMS, GML, ZAM

Drug formulation/supply -- SHO

Data collection – SES, ARB, JKA, AES, IG, MLB

Data analysis – SES, ARB, ZAM

Writing/editing -- SES, ARB, LMS, GML, ZAM

---

**Supplemental Figure 1.**
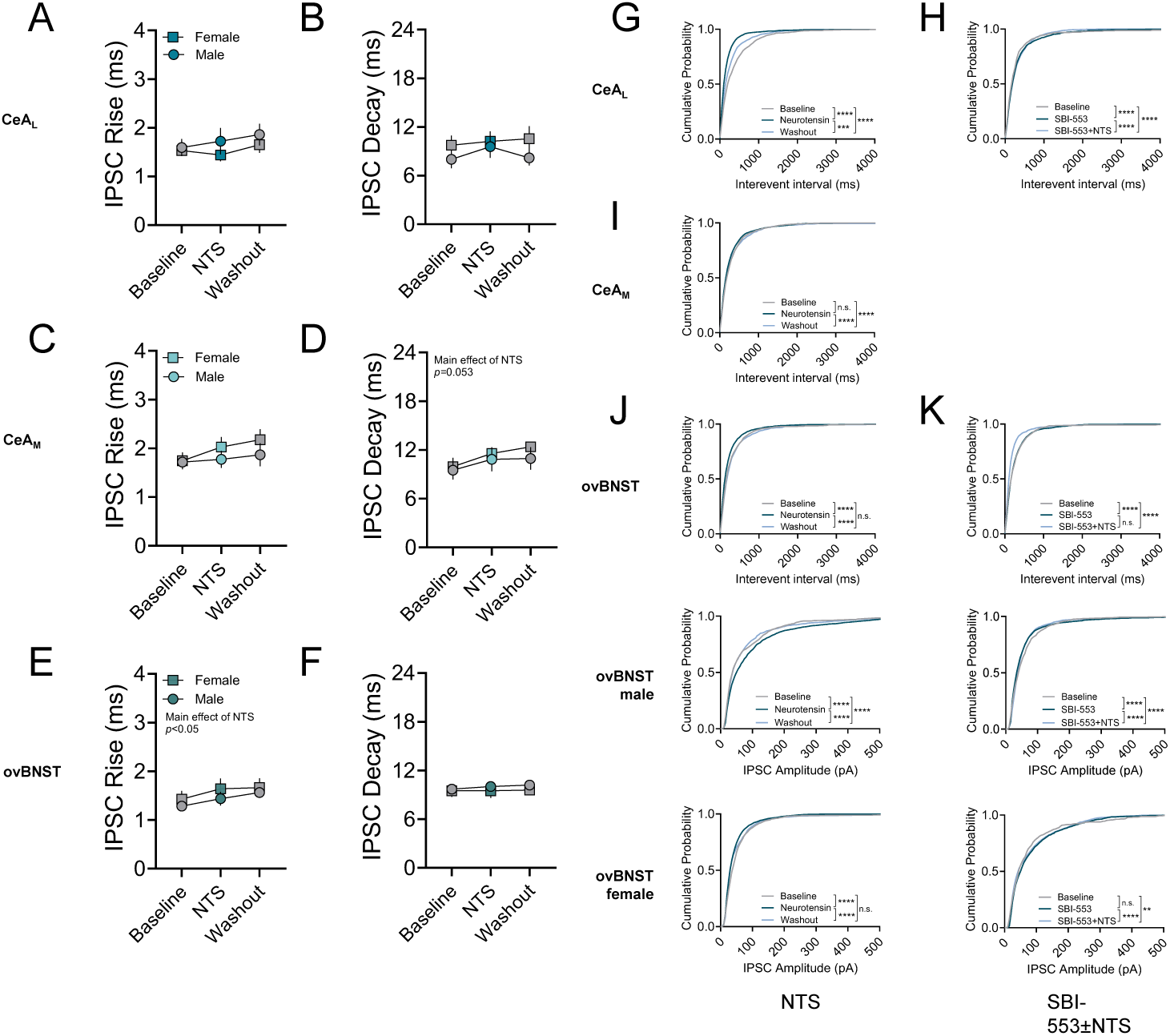
cumulative distributions

**Supplementary Fig. 2.**
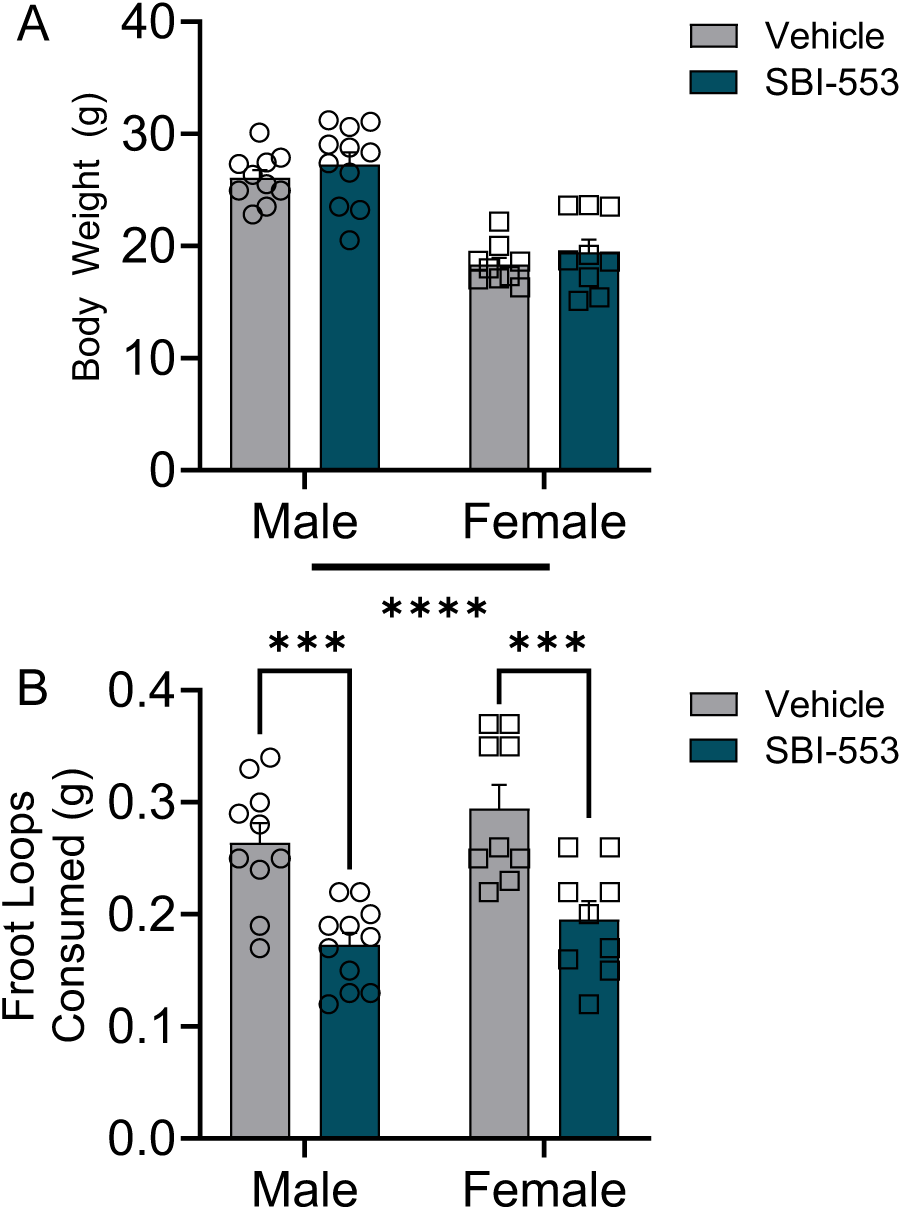
MHFA consumption sex differences are driven by bodyweight Sex differences in MHFA consumption are driven by bodyweight. **(A)** Males weighed significantly more than females but there were no differences in pre-treatment bodyweight between treatment groups. **(B)** Males and females consumed similar amounts of FL under both vehicle and SBI-553 treatment. Error bars represent average value ±SEM.

**Supplementary Fig. 3.**
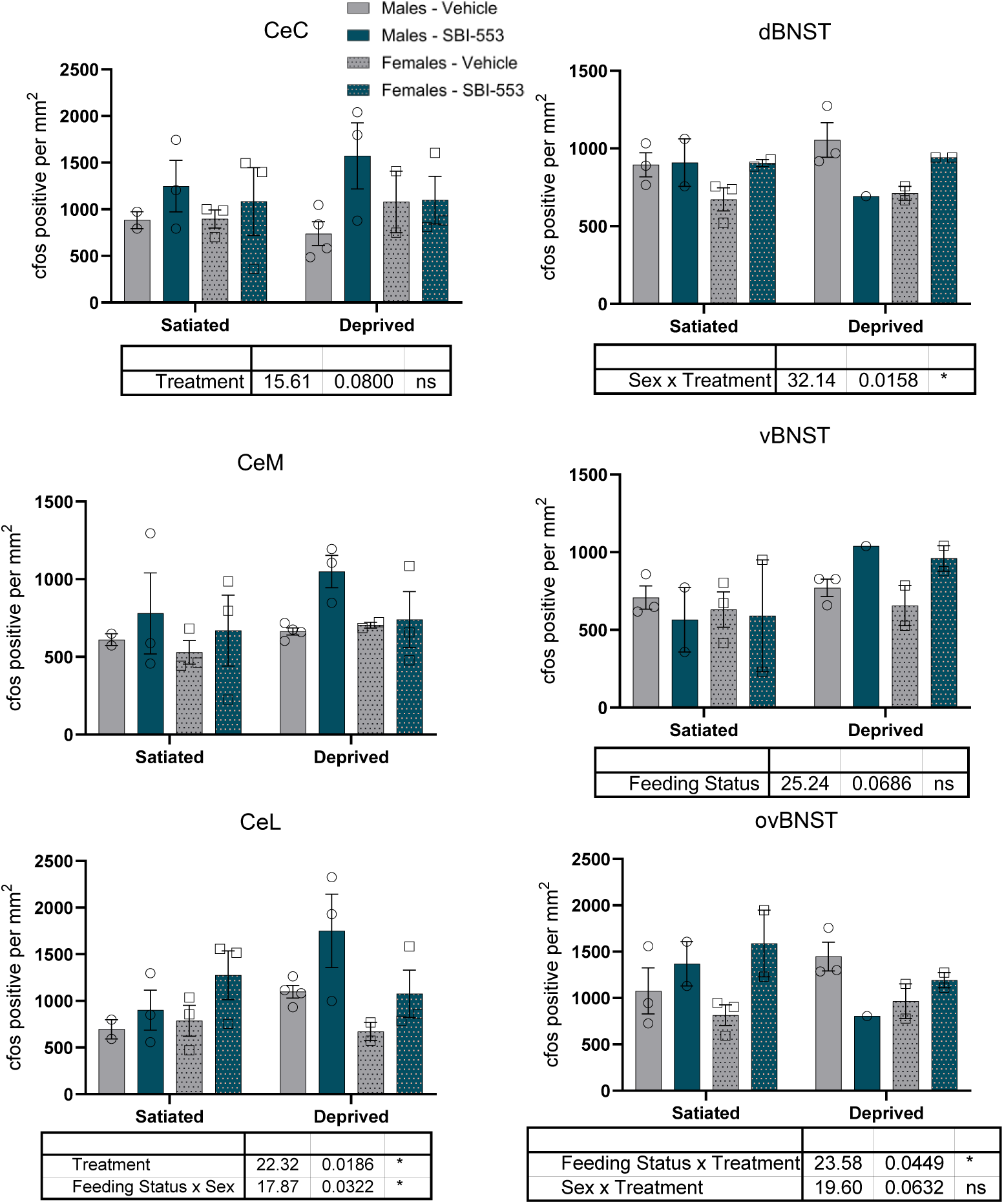
FvFHFA cFos activity by Feeding Condition, Sex, and Treatment cFos expression in the CeA and BNST separated by feeding condition, sex, and treatment. **(A-C)** cFos expression in the central amygdala and **(D-F)** BNST subregions after SBI-553 or vehicle treatment in the FvFHFA. Error bars represent average value ±SEM. Tables below each graph show ANOVA results with P<0.1 (* = P<0.05)

**Supplementary Fig. 4.**
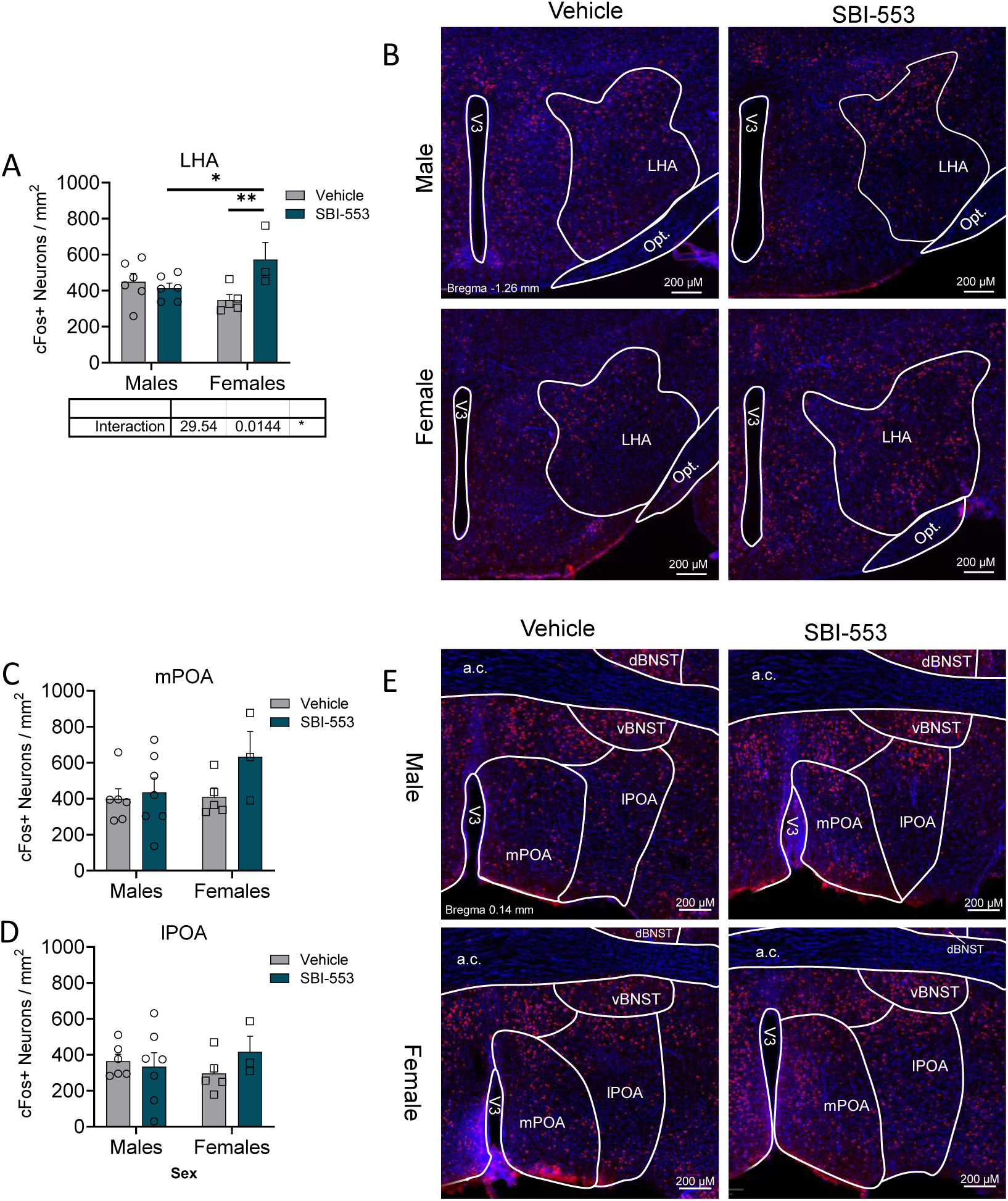
– MHFA cFos activity in mPOA and LHA Impact of SBI-553 on cFos activity in the Lateral Hypothalamus (LHA) and preoptic area (POA) after the MHFA. **(A)** Females treated with SBI-553 showed higher cFos expression compared to SBI-553 treated males or vehicle treated females. **(B)** Representative immunofluorescence images of cFos expression (red) and DAPI (blue) in the LHA. **(C)** SBI-553 had no impact in either males or females on cFos expression in the mPOA or **(D)** lPOA. **(E)** Representative immunofluorescence images of cFos expression in the POA. Error bars represent average value ±SEM. *p<0.05, **p<0.01

**Supplementary Fig. 5.**
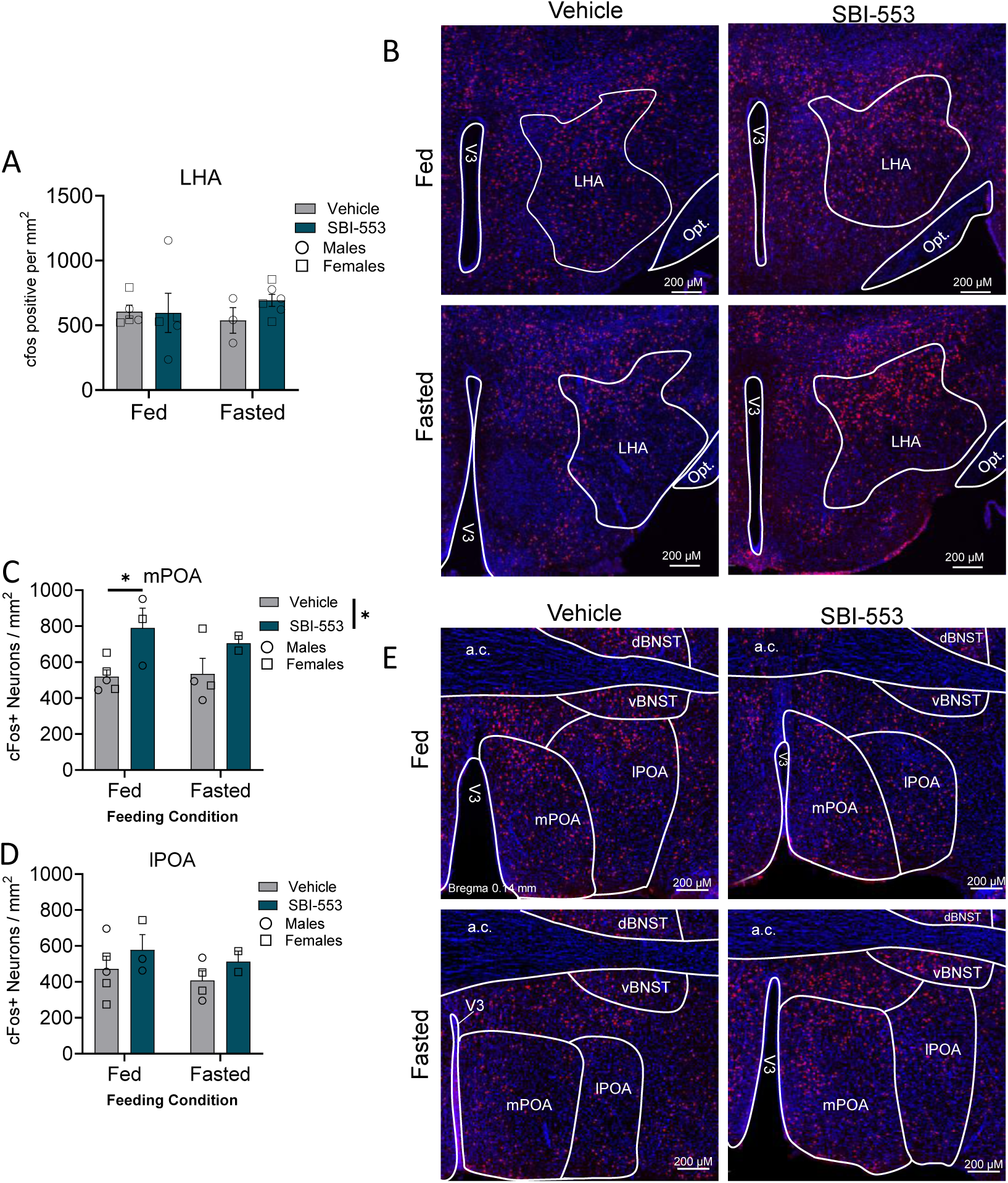
– FvFHFA cFos activity in mPOA and LHA Impact of SBI-553 on cFos activity in the Lateral Hypothalamus (LHA) and preoptic area (POA) after the FvFHFA. **(A)** Neither SBI-553 nor feeding state impacted cFos activity in the LHA. **(B)** Representative immunofluorescence images of cFos expression (red) and DAPI (blue) in the LHA. **(C)** SBI-553 significantly increased cFos expression in the mPOA **(D)** but not lPOA after the FvFHFA. **(E)** Representative immunofluorescence images of cFos expression in the POA. Error bars represent average value ±SEM. *p<0.05

**Supplemental Table 1-.**
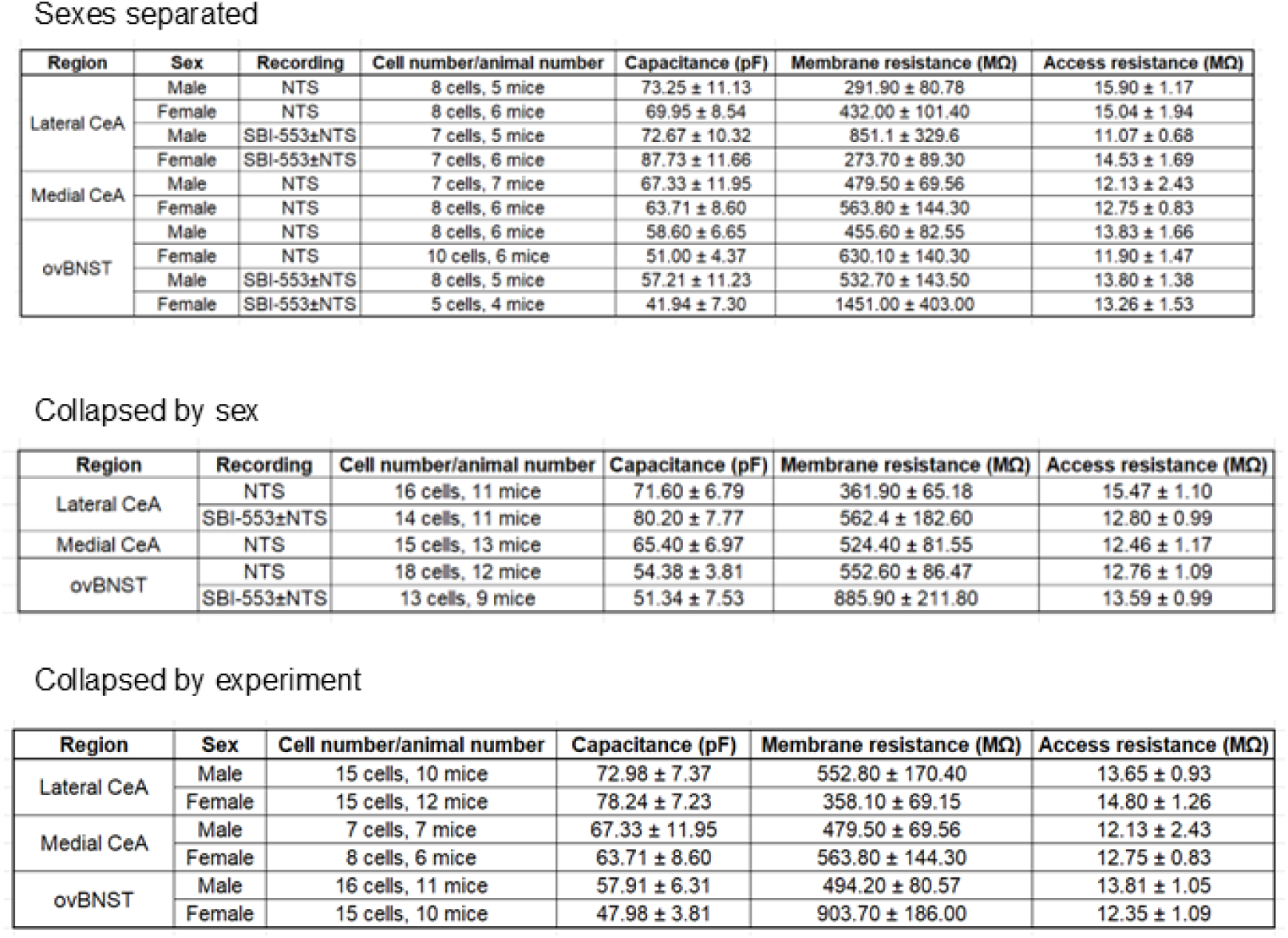
Membrane Properties for electrophysiology recordings.

**Supplemental Table 2.** – Nutritional Comparison of Chow and Froot Loops. Nutritional comparison of standard rodent chow (PicoLab Select Rodent 50 IF/6F, 5V5R) and palatable food (Froot Loops, WK Kellogg Co) used in feeding experiments.

| Nutrient | 5V5R Chow | Froot Loops |
| --- | --- | --- |
| Protein (%) | 19.0 | 5.1 |
| Fat (%) | 5.9 | 3.8 |
| Sugar (%) | 0.32 | 30.8 |
| Fiber (%) | 2.4 | 5.1 |
| Total Carbohydrate (%) | 57.6 | 87.2 |
| Energy (kcal/g)* | 3.60 | 3.85 |
| Calories from Protein (%) | 21.1 | 5.3 |
| Calories from Fat (%) | 14.8 | 9.0 |
| Calories from Carbohydrate (%) | 64.0 | 85.7 |
| Sodium (%) | 0.22 | 0.54 |

\*Energy density calculated using Atwater factors (protein × 4, fat × 9, carbohydrate × 4 kcal/g).

## Glossary

aCSF: artificial cerebral spinal fluid
BAM: biased allosteric modulator
BNST: bed nucleus of the stria terminalis
BNST^CRF^: bed nucleus of the stria terminalis, corticotropin-releasing factor neurons
BNST^NTS^: bed nucleus of the stria terminalis, neurotensin neurons
BNST^pdyn^: bed nucleus of the stria terminalis, prodynorphin neurons
CeA: central amygdala
CeAC: central amygdala, capsular subregion
CeA^CRF^: central amygdala, corticotropin-releasing factor neurons
CeAL: central amygdala, lateral subregion
CeAM: central amygdala, medial subregion
CeA^NTS^: central amygdala, neurotensin neurons
CeA^PKCδ^: central amygdala, protein kinase Cδ neurons
CeA^SOM^: central amygdala, somatostatin neurons
DAG: diacylglycerol
dBNST: bed nucleus of the stria terminalis, dorsal subregion
FL: froot loop
FvFHFA: fasted vs fed home cage feeding assay
GPCR: G-protein coupled receptor
IPSC: inhibitory postsynaptic current
LHA: lateral hypothalamus
lPOA: lateral preoptic area
LTP: long-term potentiation
mPOA: medial preoptic area
NSFT: novelty-suppressed feeding test
NTS: neurotensin
NTSR1: high affinity neurotensin 1 receptor
NTSR2: low affinity neurotensin 2 receptor
ovBNST: bed nucleus of the stria terminalis, oval nucleus
PAG: periaqueductal gray
PBN: parabrachial nucleus
PLC: phospholipase C
sIPSC: spontaneous inhibitory postsynaptic current
SN: substantia nigra
vBNST: bed nucleus of the stria terminalis, ventral subregion
VTA: ventral tegmental area

## Notes

### Competing Interest Statement

US Patent 9,868,707 relating to composition of matter for SBI-553 and its derivatives was issued to the Sanford Burnham Prebys Medical Discovery Institute (SBP) and Duke University, and US Patent 10,118,902 has been issued to SBP. Patent application US20240398806 related to the use of SBI-553 and its derivatives has been filed by Duke University. The authors have no financial interests to disclose.

### Summary of Updates

In this revision we added in new data, fixed some errors in prior analysis that were caught following the first submission, and added in addtional statistical metrics (effect size). We also expanded upon concepts in the intro, methods, results, and discussion.

